# Carbon signaling protein SbtB possesses redox-regulated apyrase activity to facilitate regulation of bicarbonate transporter SbtA

**DOI:** 10.1101/2022.05.18.492403

**Authors:** Khaled A. Selim, Michael Haffner, Reinhard Albrecht, Hongbo Zhu, Karl Forchhammer, Marcus D. Hartmann

**Author notes:** corresponding authors: K.A. Selim & M.D. Hartmann. **Authors E-mails list:** K.A. Selim. ORCID information KAS – https://orcid.org/0000-0002-2974-9483. KF – https://orcid.org/0000-0003-3199-8101. MDH - https://orcid.org/0000-0001-6937-5677.

## Abstract

The PII superfamily consists of widespread signal transduction proteins found in all domains of life. In addition to canonical PII proteins involved in C/N sensing, structurally similar PII-like proteins evolved to fulfill diverse, yet poorly understood cellular functions. In cyanobacteria, the bicarbonate transporter SbtA is expressed with the conserved PII-like protein, SbtB, to augment intracellular C_i_ levels for efficient CO_2_-fixation. We identified SbtB as a sensor of various adenine nucleotides including the second messenger nucleotides cAMP, known as carbon-status indicator, and c-di-AMP, involved in global cellular homeostasis. Moreover, many SbtB proteins possess a C-terminal extension with a disulfide bridge. We previously implied a redox-regulatory function of this extension, which we now call R-loop. Here, we reveal an unusual ATP/ADP apyrase (diphosphohydrolase) activity of SbtB that is controlled by the R-loop. We followed the sequence of the hydrolysis reactions from ATP over ADP to AMP in crystallographic snapshots and reveal the structural mechanism by which changes of the R-loop redox state modulate apyrase activity. We further gathered evidence that this redox state is controlled by thioredoxin TrxA, suggesting that it is generally linked to cellular metabolism. Finally, we present a refined model of how SbtB regulates SbtA activity, in which both the apyrase activity and its redox regulation play a central role. This highlights SbtB as a central switch-point in cyanobacterial cell physiology, integrating not only signals from the energy state (adenyl-nucleotide binding) and the carbon supply via cAMP binding, but also from the day/night status reported by the C-terminal redox-switch.

## Introduction

The proteins of the PII signal transduction superfamily are widespread in all domains of life, representing one of the most ancient and largest signaling protein families in nature (Forchhammer et al. 2022). These proteins are characterized by their highly conserved trimeric structure, consisting of a triangular core of β-sheets from the ferredoxin-like fold of the three subunits (Forchhammer & Lüddecke 2016; Wheatley et al. 2016; Selim et al. 2018, 2020a and 2021b). These subunits have three characteristic loop regions (T-, B- and C-loops) (Selim et al. 2018, 2020a and 2022; Kaczmarski et al. 2019; Liu et al. 2021; Fang et al. 2021), which are located near the inter-subunit clefts and play a major role in ligand binding and intramolecular signaling. Despite their highly conserved structure, their amino acid sequence conservation is often low, implying that the PII superfamily also comprises members that are involved in the regulation of cellular activities that differ markedly from those controlled by canonical PII proteins (Forchhammer & Lüddecke 2016; Wheatley et al 2016; Selim et al. 2018, 2021b and 2022). In cyanobacteria, in addition to the canonical PII protein involved in N/C sensing (Forchhammer & Selim 2020; Forchhammer et al. 2022), the PII-like protein SbtB evolved to regulate the cyanobacterial carbon-concentrating mechanism (CCM) (Selim et al. 2018).

Cyanobacteria use the CCM to cope with limiting CO_2_ levels, augmenting intracellular levels of inorganic carbon (hereinafter C_i_; referring to bicarbonate and CO_2_) and providing the major carbon fixing enzyme, ribulose 1,5-bisphosphate carboxylase/oxygenase (RubisCO), with CO_2_ to ensure efficient carboxylation of ribulose 1,5-bisphosphate and repress its oxygenation reaction (Forchhammer & Selim 2020). Cyanobacterial CCM comprises several CO_2_ and HCO_3_^-^ uptake systems. The high affinity sodium-dependent bicarbonate transporter SbtA is one component of the CCM system and is highly expressed under C_i_ limitation together with its downstream gene encoding for SbtB. Previous works demonstrated that SbtA is the primary target of SbtB (Du et al. 2014; Selim et al. 2018; Liu et al. 2021; Fang et al. 2021). In our previous work, we provided first structural, biochemical and physiological characterization of this unique PII-like protein from *Synechocystis* (*Sc*SbtB) (Selim et al. 2018; Mantovani et al. 2022). The binding properties of *Sc*SbtB turned out to be unique among the so-far characterized members of the PII superfamily, as it can bind a variety of adenine nucleotides (ATP, ADP, and AMP), including the second messenger cyclic AMP (cAMP). Recently, we additionally reported the binding of the second messenger c-di-AMP to *Sc*SbtB, which plays a key role in the regulation of diurnal and carbon metabolism via controlling glycogen synthesis (Selim et al. 2021a; Mantovani et al. 2022). We could demonstrate that c-di-AMP binding promoted the interaction of SbtB with the glycogen-branching enzyme GlgB, identifying the latter as another important target of SbtB (Selim et al. 2021a).

Of the different adenyl nucleotides, *Sc*SbtB exhibited the highest affinity for cAMP and c-di-AMP, followed by ADP, ATP and then AMP. All of them bind in the canonical binding sites at the inter-subunit clefts and the binding modes of AMP (PDB: 5O3R), cAMP (PDB: 5ORQ) and c-di-AMP (PDB: 7OBJ) were disclosed in crystal structures. Intriguingly, although AMP and cAMP were found to convey opposite signals, a comparison of the *Sc*SbtB:AMP and *Sc*SbtB:cAMP complexes did not reveal any conformational differences on the flexible surface-exposed T-loop, the canonical protein interaction module of PII proteins. In contrast, the *Sc*SbtB:c-di-AMP complex revealed a structural rearrangement of this loop in response to c-di-AMP binding.

It is generally evident that SbtB exerts its regulatory functions on SbtA and GlgB through the differential binding of these adenine nucleotide effector molecules, which may induce conformational changes on the T-loop, in analogy to canonical PII proteins (Forchhammer & Selim 2020; Forchhammer et al. 2022). This is underlined in recent studies revealing the structure of the SbtB:SbtA complex (Fang et al., 2021; Liu et al. 2021), showing a symmetric binding of the trimeric SbtB protein to a trimeric SbtA structure in which the T-loops of SbtB lock partially the C_i_ channels of SbtA (Fang et al. 2021; Liu et al. 2021). This complex is stabilized by the AMP-bound state of SbtB, but sterically incompatible with cAMP and ATP binding (Fang et al. 2021; Liu et al. 2021). However, the structural details of how SbtB regulates other targets, like GlgB, remain elusive.

All available data imply that SbtB regulates the activity of its targets in response to the concentrations of intracellular adenyl-nucleotides and the redox or photosynthetic state of the cell (i.e. day-night transition). In this context, a further striking feature of *Sc*SbtB is a C-terminal extension that contains a highly conserved CGPxGC motif, which is widespread among SbtB proteins (Selim et al. 2018). This extension forms a small hairpin loop structure, in which a disulfide bond is found between the two cysteines C105 and C110 (Selim et al. 2018). We hypothesized that this extension might be involved in a redox-sensory function that may read out the redox or photosynthetic state of the cell and integrate this information into the regulation of its targets; in the following we will call this extension R-loop (for **<u>R</u>**edox-regulated loop). In this study, we used structural and biochemical approaches to reveal an intricate interplay of nucleotide binding and the redox state of the R-loop. We found that SbtB slowly hydrolyzes the adenine nucleotides ATP and ADP to AMP, show that this activity is redox-regulated via the R-loop and suggest that it serves to coordinate SbtA:SbtB complex formation in response to the day-night cycle.

## Results

### SbtB has ATP diphosphohydrolase (apyrase) activity

In our initial structural characterization of SbtB (Selim et al. 2018), we were able to solve crystal structures of *Sc*SbtB in apo-state as well as in complex with AMP and cAMP, which all crystallized in the same trigonal crystal form in space group P3_2_, irrespective of the ligandation state. *Sc*SbtB resembles the canonical trimeric PII architecture, but has a peculiar CGPxGC motif at its C-terminus, which is conserved in many but not all SbtB homologs in cyanobacteria (Fig. S1) (Selim et al. 2018). In this motif, which we term the R-loop (for **<u>R</u>**edox-regulated loop), the two cysteines were found to form a disulfide bridge (PDBs: 5O3Q and 5ORQ). Moreover, the carboxyl terminus of the last cysteine can form a hydrogen bond with the amino terminus, which appeared to stabilize the whole R-loop assembly. The adenine nucleotides, either AMP or cAMP, were found to form the canonical interactions in the inter-subunit clefts, with their phosphate groups mostly solvent-exposed and the T-loop largely disordered. To our surprise, co-crystallization attempts with ADP (Selim et al. 2018) and also ATP, which also yielded the same trigonal crystal form, contained only AMP in all three binding sites, suggesting that the nucleotides were hydrolyzed during the time course of the crystallization experiment. To find out if the purified recombinant Strep-tagged *Sc*SbtB preparation contained a contaminating ATPase/ADPase activity, the *Sc*SbtB preparation was analyzed by tandem-mass spectrometry (MS/MS) (Fig. S2 and proteomic dataset 1). As no contamination with potential phosphate hydrolase activity could be detected, we speculated that *Sc*SbtB might itself hydrolyze the adenine nucleotides, similar to what has been reported for canonical PII proteins (Radchenko et al. 2013). Therefore, we assayed for phosphorylase activity using ATP and ADP as substrates and confirmed slow ATP and ADP hydrolysis activity by *Sc*SbtB, which was 5-fold accelerated in presence of Mg^2+^ (Fig. 1 and Fig. S3). A very similar activity was obtained using a different recombinant SbtB protein, derived from filamentous cyanobacterium *Nostoc* sp. PCC 7120, indicating that the hydrolysis of adenine nucleotides is a common trait among cyanobacterial SbtB proteins. To reveal the specificity of metal ions on phosphate release, various divalent metal ions at 5 mM concentration were added to the assay (Fig. S3), revealing that also Mn^2+^ and Co^2+^ stimulate phosphate release in addition to Mg^2+^, whereas all other metals were ineffective. Since Co^2+^ is toxic for cyanobacteria and cannot be found in excess inside the cell (Selim & Haffner 2020), we concluded that Mg^2+^ and Mn^2+^ are most likely the physiologically relevant metals used by SbtB.

**Fig. 1.**
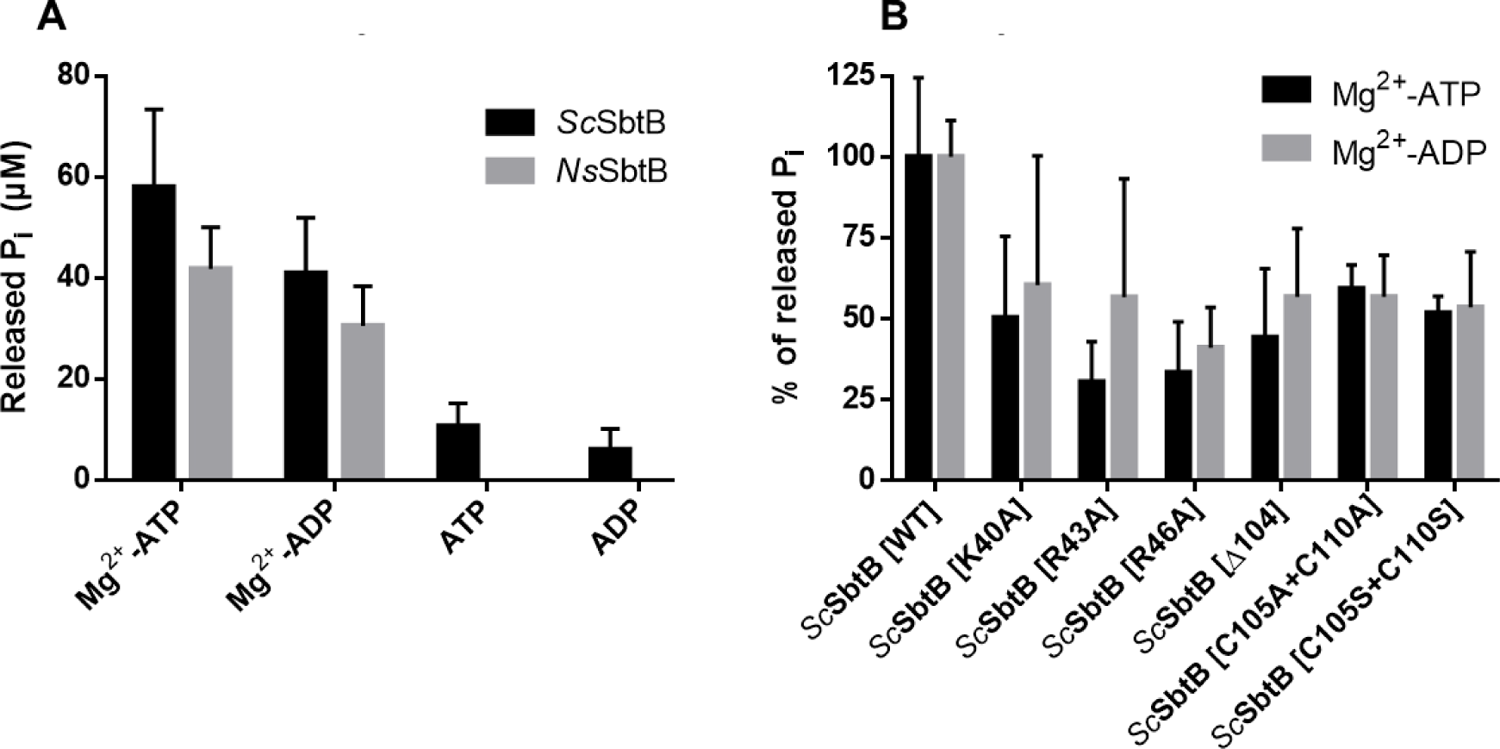
Apyrase activity of SbtB via phosphate release assay. (A) Phosphate release assay for *Sc*SbtB and *Ns*SbtB in presence and absence of Mg^2+^ ions. The released inorganic phosphate (P_i_) is shown in µM. (B) Relative phosphate release activity of different SbtB variants compared to wildtype *Sc*SbtB (100%), as indicated. Values are means ± SD; *n* = 5 to 6 independent measurements of different SbtB purifications.

### Following ATP/ADP hydrolysis in crystallographic snapshots

As hydrolysis appeared to be a slow process (Fig. S3), we tried to study it structurally by performing crystal soaking experiments. To this aim, we incubated the P3_2_ crystals of apo-*Sc*SbtB, in which the R-loop is folded via the disulfide bridge, in their crystallization solution supplemented with either ADP or ATP for different time spans. These crystals yielded several datasets with resolutions better than 2.5 Å and well-defined electron density for the unambiguous identification of β- and γ-phosphate groups of ADP and ATP molecules (Fig. S4). Together, the different structures illustrate different time points of a potential hydrolysis reaction from bound ATP over ADP to AMP: While a short (2 h) ATP soak shows clear electron density for ATP in all three binding sites, ATP is found to be hydrolyzed to ADP in a long (overnight) ATP soak, and ADP is found mostly hydrolyzed to AMP in a (4 h) ADP soak. Compared to the SbtB:AMP structure, the complexes with ADP and ATP show additional interactions between their β- and γ-phosphate groups and the protein, which is most pronounced in the short ATP soak (Fig. 2).

**Fig 2.**
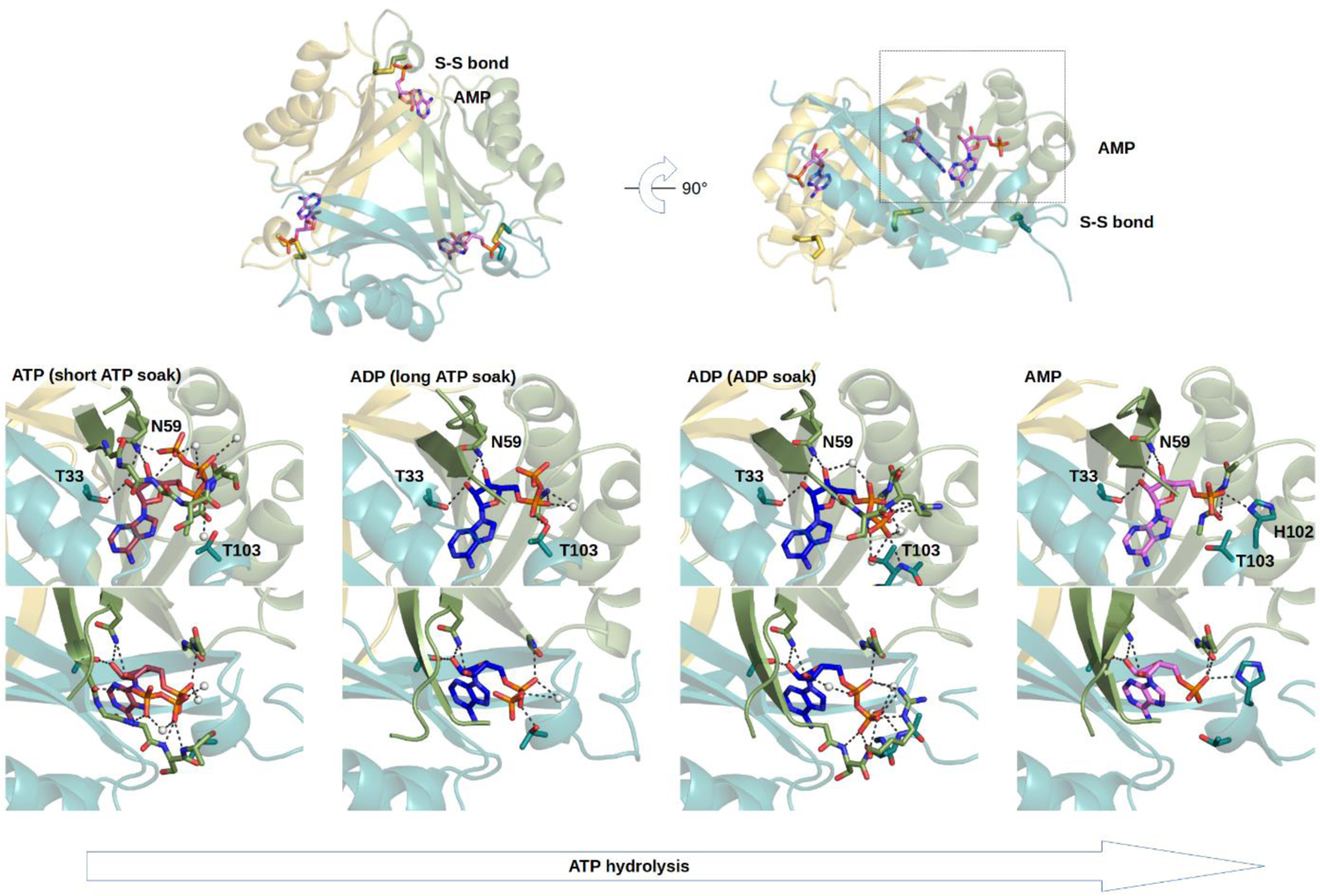
Overall structure of SbtB and sequence of ATP/ADP hydrolysis with an oxidized R-loop. Top: Trimeric structure of the SbtB:AMP complex. The nucleotides are bound between the subunits, the T-loops are largely unfolded and indicated by dashed lines. The disulfide bonds in the oxidized R-loop are indicated in stick representation. Bottom: Structural snapshots along the sequence of hydrolysis when the R-loop is oxidized. After the short ATP soak, fully occupied ATP is found in all three binding sites. After prolonged soaking times, ATP is found to be hydrolyzed to ADP, with the β-phosphate in the same orientation as in the ATP-bound structure. In the ADP soak, the β-phosphate is found in a different orientation, which might represent the orientation that needs to be assumed for ADP hydrolysis. Finally, the AMP-bound structure represents the end of the full hydrolysis sequence, which can be obtained by co-crystallization with either AMP, ADP or ATP. Na^+^ ion forming the nucleotide binding pocket is shown as a gray sphere. Electron densities for the individual ligands are shown in (Fig. S4).

In the short ATP soak, the three chains of trimeric *Sc*SbtB differ noticeably in the area of the effector molecule binding cleft, where the T-loop is structured to different extents. Compared to the AMP-bound structure (PDB: 5O3R), where the T-loop is disordered after residue G41, the base of the T-loop is resolved until R43 in one, and the T-loop is resolved until S48 in another chain (chain C), where it wraps around the phosphate groups. ATP is found in an unusually strained conformation, in which one of the γ-phosphate oxygens (Oγ1) is forming a 2.6 Å intramolecular hydrogen bond to the 3’-hydroxyl group of the ribose moiety. The Oγ2 oxygen is forming a hydrogen bond with the G41 backbone nitrogen, and in chain C, Oγ3 and Oγ1 are forming salt bridges with the R46 guanidino group. Also, in chain C, the β-phosphate forms hydrogen bonds with the N44, V45 and R46 backbone nitrogen atoms, while the α-phosphate forms hydrogen bonds with the S42, R43 and G89 backbone nitrogen atoms and a salt bridge with the R43 guanidino group. In two of the chains, the coordination of ATP is completed with a metal ion bridging the β- and γ-phosphates, which is further coordinated by N44 in chain C. Based on the electron density and coordination distances of about 2.4 Å, we interpreted this metal as a sodium ion from the crystallization buffer, mimicking a potentially physiologically relevant Mg^2+^ ion. Of note, the described T-loop conformation in chain C has a weaker electron density than the rest of the protein, while the residues preceding the R-loop have a weaker electron density as well. Closer inspection reveals that T-loop residues S42 and R43 are in too-close contact to R-loop residues G101, H102 and T103, such that both stretches cannot assume the described conformation simultaneously. Accordingly, these residues were modelled with partial occupancy, reflecting the overall weaker electron density. Taken together, we observe that the T-loop can wrap around and coordinate the phosphates of ATP, but this conformation is conflicting with the folded R-loop.

In the long ATP soak, we find that ATP was hydrolyzed to ADP in two of the chains, and the start of the T-loop is only structured until G41 in all three chains. The β-phosphates of the ADP molecules are found in the same orientation as in the ATP molecules prior to hydrolysis, but do not form additional interactions with SbtB as the T-loop is unstructured, while the R-loop is ordered as in the other structures. In the ADP soak, however, we find ADP in another conformation (Fig. 2). While it is already converted to AMP in two chains, in the third chain, it has the β-phosphate relocated towards the R-loop, where it interacts with two additionally folded T-loop residues, with the S42 backbone nitrogen and the R43 guanidino group. While this could possibly be an artifact of the soaking procedure, it is conceivable that this conformational switch of the ADP molecule is a necessary part in consecutive steps of the hydrolysis reactions from ATP to ADP and from ADP to AMP. In the light of these results, we concluded that SbtB might display redox-dependent ATP/ADP diphosphohydrolase (Apyrase) activity. As the folded (oxidized) R-loop is conflicting with a conformation in which the T-loop is wrapping around the ATP phosphates, we assumed that its function might be to promote hydrolysis, while a reduced and thus unfolded R-loop might allow a tighter binding and thus stabilization of ATP.

### T-loop Arginines and oxidized R-loop are critical for ATP hydrolysis

The observed nucleotide binding modes, the stepwise hydrolysis of ATP and the apparent incompatibility of T-loop folding with the oxidized R-loop inspired a number of experiments, for which we constructed two sets of mutants. The first set comprises point mutants in the T-loop, in which we substituted either R43, R46 or K40 by alanine, which are involved in the coordination of the β- and γ-phosphates and the metal ion coordinating these phosphates in the short ATP soak (PDB: 7R2Y), respectively. The second set was inspired by the apparent incompatibility of T-loop- and R-loop-folding. It comprises variants in which we mutated both R-loop cysteines (C105 and C110) to either alanine or serine (C105A+C110A or C105S+C110S) such that the R-loop should not be able to assume its folded structure due to the lack of the disulfide bond, and a variant in which we truncated the R-loop from position 104 on (Δ104). With both sets of variants, we performed phosphate release assays, and in fact, all variants showed reduced phosphate release activity, both with Mg^2+^-ATP and Mg^2+^-ADP as substrate (Fig. 1).

These results have two implications. Firstly, they show that R43, R46 and K40 are not only important for the binding of ADP and ATP, but also for their breakdown. Secondly, while the folded R-loop supports this breakdown, hydrolysis is efficiently inhibited when the R-loop is not folded or absent, which is especially interesting in the light of our previous hypothesis that the R-loop might function as a redox switch (Selim et al. 2018).

### The T-loop can adopt an ATP-protecting conformation in absence of a folded R-loop

As our initial crystal structures already suggested that a structured R-loop is incompatible with the folding of the T-loop, we decided to perform structural studies on *Sc*SbtB variants which are either lacking the R-loop (Δ104), or mimicking its reduced state using the alanine or serine substitution variants (C105A+C110A or C105S+C110S). For simplicity, we will refer to both variants as SbtB^defR^ (for deficient in/defunctional R-loop), where applicable. With those variants, we performed co-crystallization trials with ADP and ATP, which all yielded the same crystal form in space group P4_1_. For ADP, the best dataset was obtained with the Δ104 variant, which was scaled to 1.8 Å resolution (SbtB^defR^:ADP), while the best dataset with ATP was obtained with the C105A+C110A variant, which was scaled to 1.5 Å resolution (SbtB^defR^:ATP).

The structures have one SbtB trimer in the asymmetric unit and show unambiguous electron density for their respective nucleotide in all three binding sites (Fig. S4). Strikingly, the two structures are very similar to the ADP- and ATP-bound structures recently reported for SbtB from *Cyanobium* sp. PCC7001, which belongs to a group of SbtB proteins lacking the R-loop extension (Figs. S1 and S5, Kaczmarski et al. 2019). In all chains of the SbtB^defR^:ATP and SbtB^defR^:ADP structures, the base of the T-loop assumes a similar conformation as the one observed in one chain of the short ATP soak, which was incompatible with the folded R-loop (see above; Fig. S6). However, besides numerous similarities, the ADP- and ATP-bound structures also show peculiar differences in their nucleotide binding modes and T-loop conformations (Fig. 3).

**Fig 3.**
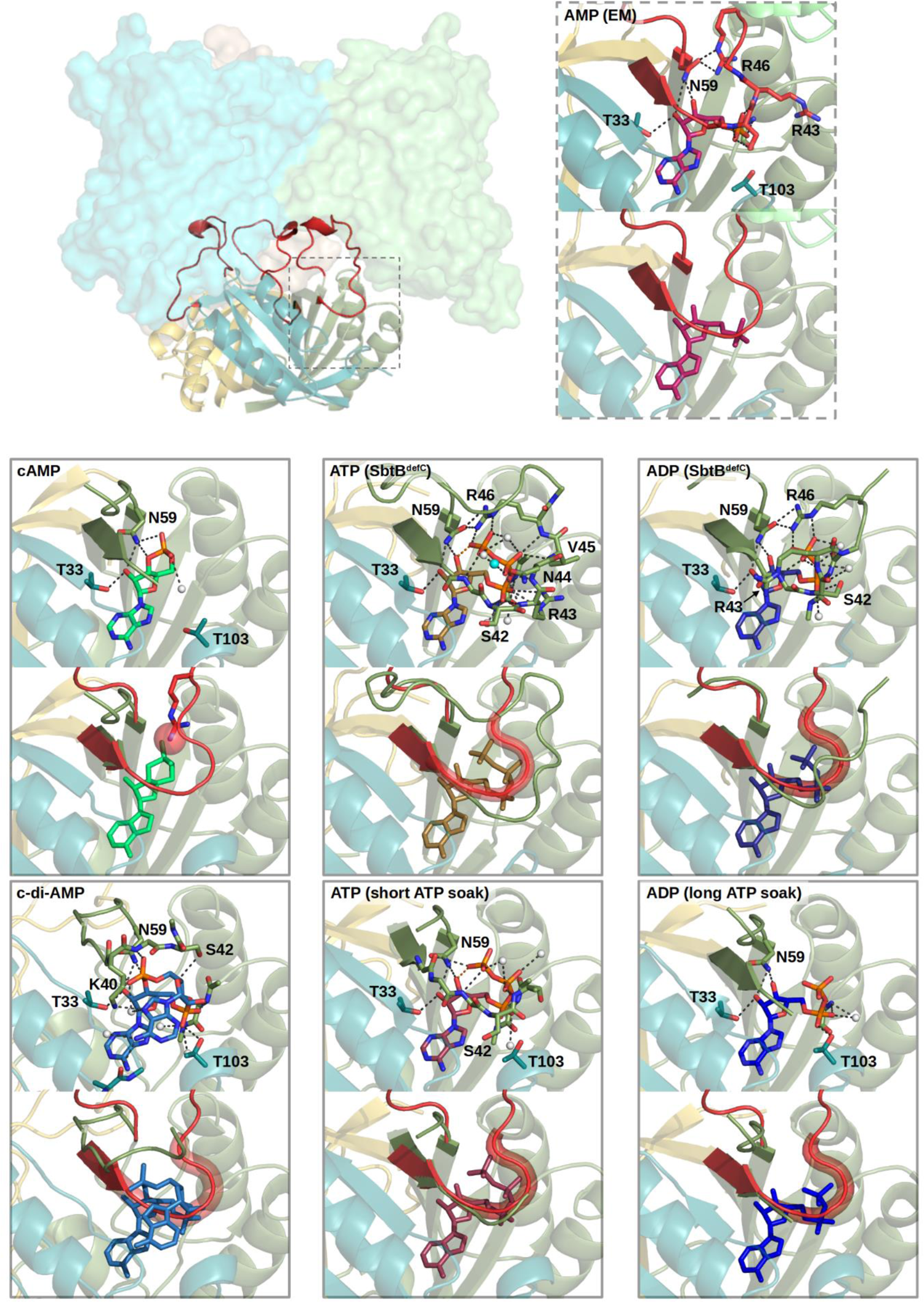
Complex structures illustrating how the SbtA:SbtB association is regulated by nucleotide binding. In the SbtA:SbtB complex, the SbtB T-loops are inserted into cavities of the individual SbtA subunits, stabilized by AMP molecules. This particular T-loop conformation is colored red, and anchored by an interaction between R46 and N59. In the complex structures SbtB:cAMP, SbtB:c-di-AMP, SbtB^defR^:ATP, SbtB:ATP, SbtB^defR^:ADP and SbtB:ADP, this T-loop conformation is prevented in different ways. For each complex, both the detailed nucleotide binding modes and the superposition to the T-loop in SbtA-bound conformation (red) are illustrated. In the superpositions, the areas in which the SbtA-bound T-loop conformation would clash with the bound nucleotide are highlighted with a red glow. In the cAMP-bound state, the cAMP phosphate would preclude R46 from assuming its anchoring role, while in the c-di-AMP state, the base of the T-loop is found in an entirely different conformation. For the ATP and ADP state with an oxidized R-loop (short and long ATP soaks), the phosphate groups of the nucleotides would obstruct the T-loop path. The same is the case for the ATP and ADP state with a reduced R-loop (mimicked in SbtB^defR^), where the T-loop is partly or completely folded around the phosphate groups: In SbtB^defR^:ATP the T-loop is fully ordered and wrapped around the phosphates, forming a sophisticated network of interactions, which is completely different form the T-loop conformation in the StbA-bound state. In analogy to the latter, R46 is also interacting with N59, but in a completely different orientation, additionally coordinating the γ-phosphate (See also Fig. S5). In SbtB^defR^:ADP, the interaction network is similar but less complete. Strikingly, R46 is in a similar orientation as in the reduced ATP state, but coordinating the β-phosphate instead of the γ-phosphate group.

In the SbtB^defR^:ATP structure, the R-loop is largely disordered (Fig. S6) and the T-loop is entirely folded in two chains, and in all three chains, all nucleotide interactions are formed as described for the most complete chain (C) in the short ATP soak, including the sodium ion with a well-resolved complete octahedral coordination sphere (Fig. 3). As this T-loop conformation wraps around the phosphate groups of the nucleotides to potentially prevent their hydrolysis, we call this the “protecting state”. Of note, the functionally important R46, which is coordinating the γ-phosphate, is additionally forming a hydrogen bond with N59, which was also observed in the short ATP soak. N59 generally forms hydrogen bonds with the hydroxyl groups of the ribose of all bound nucleotides, but in the protecting state, it has its side chain flipped to additionally bind to R46, which presumably stabilizes the T-loop in this conformation.

In the SbtB^defR^:ADP structure, the T-loop is not fully resolved, but in two chains, it is found in a conformation similar to the ATP-protecting state. ADP is found in a similar orientation as in the post-hydrolysis state, with the β-phosphate in a similar location as the ATP γ-phosphate. Additionally, R46 is found in essentially the same conformation as in the ATP-protecting state, but it is now interacting with the β-phosphate instead of the γ-phosphate, and it forms the same stabilizing hydrogen bond with N59 as in the ATP-protecting state. However, there are significant conformational differences in the T-loop residues preceding R46, which interact with both the α- and β-phosphate in the ATP-protecting state. Most prominently, R43 in the SbtB^defR^:ADP complex is found in a very different location, where it is not interacting with the α-phosphate but with the β-phosphate. In this overall conformation, the path of the T-loop backbone, most importantly that of S42 and R43, would not necessarily clash with a folded R-loop (Fig. S6). Therefore, we assume that this conformation is also possible in wildtype *Sc*SbtB when the R-loop is folded, although it was not captured in our initial crystallization attempts.

### Structural basis for adenine nucleotide-dependent SbtA:SbtB interaction

With our previous and current crystallographic analyses, we have delineated the binding modes of the nucleotides cAMP, AMP, ADP, ATP and also c-di-AMP, and could rationalize the influence of the R-loop on the stepwise hydrolysis of ATP to AMP. In a next step, we wanted to uncover how the binding of these nucleotides influences the complex formation between SbtB and its major target SbtA. Therefore, we compared the different conformational states of the individual SbtB:nucleotide complexes to the recently reported structure of the SbtB:AMP:SbtA complex (Fig. 3). Within this complex, the T-loops are inserted into the individual SbtA subunits, forming an extensive interface, which is stabilized by the binding of AMP: At the base of the T-loop, the AMP phosphate group is coordinated by the S42 and R43 backbone nitrogens, promoting a sharp turn of the T-loop towards SbtA. Further, considering the structures of the different SbtB complexes, an interaction network between R43, R46 and N59 is of special interest. In the SbtB:AMP:SbtA complex, the R46 guanidino group forms similar hydrogen bonds with N59 as in the SbtB^defR^:ATP and SbtB^defR^:ADP structures, just that the guanidino group is in a different orientation, as the R46 backbone is not near the phosphate but buried in the interface to SbtA; on its opposite side, the R46 guanidino group forms hydrogen bonds to the R43 backbone oxygen, completing the network of interactions between these three signature residues (Fig. S5). Consequently, R46 and N59 have dual roles, as they can either stabilize the ATP-protecting state or the SbtA-bound conformation, while R43 is found in a number of different interaction modes depending on the complexation state. Although the different SbtB:nucleotide binding modes revealed a number of structural differences, they have one feature in common: All but the AMP-bound state are incompatible with the T-loop conformation required to adopt the SbtB:AMP:SbtA complex, which is illustrated in (Fig. 3). For cyclic nucleotides, bound cAMP prevents the formation of the R43-R46-N59 network, as it would clash with the cAMP phosphate group, while bound c-di-AMP completely obstructs the folding of the base of the T-loop in the required conformation, independent of the redox state of the R-loop. For the binding of ADP and ATP, the mode of obstruction depends on the R-loop. When the R-loop is folded (oxidized; Fig. S6), the β- and γ-phosphates collide with the necessary path of the T-loop (Fig. 3). With an unfolded (reduced) R-loop (Fig. S6), the additional wrapping of the T-loop around the phosphates further prevents the formation of the SbtB:SbtA interface, which is most apparent when looking at the different location and orientation of R46 (Fig. 3).

### SbtB interacts with TrxA

As the R-loop was apparently oxidized in all our studies with the wild-type protein, we asked how it can be reduced. To this aim, we first tested the influence of strong chemical reducing agents, performing crystallization trials with *Sc*SbtB in presence of TCEP and testing for apyrase activity after overnight dialysis under reducing conditions and treating the protein with 1 mM TCEP. In both experiments, the reducing agent had no apparent effect, as we obtained the same crystals in space group P3_2_ as for untreated *Sc*SbtB with a folded R-loop and the apyrase activity was not affected (Fig. S3). Consequently, we concluded that the reduction may be catalyzed enzymatically by a disulfide oxidoreductase in *Synechocystis*. Notably, we identified the *E. coli* thioredoxin (TrxA) as one of highly enriched proteins which coeluted with the strep-tagged SbtB (Fig. S2 and proteomic dataset 1). As a possible candidate, we selected the *Synechocystis* thioredoxin TrxA (encoded by *slr0623*), the only essential and most abundant thioredoxin in *Synechocystis* and all oxygenic photoautotrophs (Mallén-Ponce et al. 2021). Importantly, TrxA is also responding to day-night changes, which is known to influence the redox state of the cell (Pérez-Pérez et al. 2009). To test our assumption and examine a possible SbtB-TrxA interaction, we performed an interaction assay using the bacterial adenylate cyclase two-hybrid (BACTH) system, which we previously established for SbtB (Selim et al. 2021a). The BACTH assay depends on the reconstitution of an active adenylate cyclase (Cya) upon positive interaction of the proteins of interest fused either to the T25 and T18 subunits of Cya, which can be perceived by color change of the growing colonies to either blue or red on X-Gal or MacConkey reporter plates. Here, we fused the T25 subunit of Cya either N- or C-terminally to SbtB, while the T18 subunit of Cya was fused also N- or C-terminally to TrxA (encoded by *slr0623*) (Fig. S7). Based on our previous identification of glycogen-branching enzyme (GlgB, *sll0158*) as a target of SbtB (Selim et al. 2021a), we used T18-GlgB fusion or the leucine zipper interaction as positive controls, while the T25-SbtB fusion with an empty pUT18 vector or T18-GlgC (encoding glucose-1-phosphate adenylyltransferase by *slr1176*) were used as negative control. For three independent colonies, a clear interaction was observed only between the N-terminally tagged T25-SbtB and the C-terminally tagged T18-TrxA tagged on both of X-Gal or MacConkey plates (Fig. S7). However, only on X-Gal plates, a weak interaction was observed between the C-terminally tagged T25-SbtB and again the C-terminally tagged T18-TrxA, whereas no interaction was observed using the N-terminally tagged T18-TrxA under all tested conditions (Fig. S7). Next, we checked if the N-terminal T25-SbtB-Δ104 and the T25-SbtB-[C105A+C110A] fusions would interact with C-terminally tagged T18-TrxA. As expected, deletion of the R-loop abolished the interaction with TrxA, while the T25-SbtB-[C105A+C110A] fusion interacted weakly with TrxA. This result strongly supports our assumption that SbtB is redox-regulated via an oxidoreductase, potentially TrxA, and that the R-loop is required for the direct interaction with the oxidoreductase.

To further validate our BACTH results by another independent method, we checked for a direct physical interaction using pulldown experiments by immobilizing either recombinant streptagged *Sc*SbtB or His_6_-tagged TrxA protein on streptavidin or Ni^2+^ magnetic beads, respectively, and incubating the immobilized-protein with the other partner, followed by successive washes to remove unbound protein. As both SbtB and TrxA have a similar calculated molecular weight of 12.5-13.5 kDa, which obstructs a clear separation on SDS-PAGE gels, TrxA and SbtB were identified by immunoblotting using α-poly-His and α-strep antibodies. When SbtB was immobilized on streptavidin beads, coelution of TrxA with SbtB could be detected (Fig. S7), and *vice versa*, when TrxA was immobilized on Ni^2+^-NTA magnetic beads, wildtype SbtB was detected in the elution fraction of TrxA (Fig. S7). Remarkably, the SbtB C105A+C110A variant showed only very weak interaction with TrxA (Fig. S7), in agreement with the BACTH data, supporting our conclusion for the specificity of the TrxA-SbtB interaction to break the disulfide bond between the R-loop cysteines.

## Discussion

The unexpected apyrase activity that we unveiled for *Sc*SbtB is unusual and has a number of peculiarities and implications. While common apyrases typically hydrolyze all types of nucleotide triphosphates and diphosphates to their monophosphates, *Sc*SbtB is highly specific for adenosine nucleotides (Selim et al. 2018 and 2021a; Kaczmarski et al. 2019). This preference is due to the very specific recognition of the adenine moiety in a binding pocket that is conserved in many other proteins of the PII-like family (Forchhammer et al. 2022; Forchhammer & Lüddecke 2016). Intriguingly, a recent investigation of an SbtB-like protein known as carboxysome-associated PII protein (CPII) revealed that ADP could be hydrolyzed to AMP (Wheatley et al. 2016), although the authors could not elaborate whether or not ADP was catalytically hydrolyzed by CPII. These indications suggest that SbtB proteins generally have the ability to hydrolyze adenine nucleotides to reach to a thermodynamically stable SbtB:AMP state. Moreover, it was previously shown that canonical PII proteins possess very weak *in vitro* ATPase activity (Radchenko et al. 2013; Truan et al. 2014; Gruswitz et al. 2007), although the physiological significance of this activity remains unclear (Lüddecke & Forchhammer 2015). Considering the wide distribution and low sequence conservation between canonical PII and SbtB proteins, the fact that both show ATPase activity supports our previous assumption that the proteins of PII superfamily may have emerged from an ancestral nucleotide hydrolase (Forchhammer & Lüddecke 2016). In our structural analysis, we failed to identify residues that could serve as a general base for the deprotonation of a water molecule for the nucleophilic attack on the phosphate groups, that would be conserved among SbtB proteins. The absence of a dedicated general base might explain the slow turnover of the hydrolysis reaction, which might be facilitated by the strained conformation of the bound ATP molecule. As the apyrase activity generally leads to the AMP-bound state, in which SbtB can bind and potentially occlude the SbtA channels, it seems to generally drive the SbtA-SbtB system towards the closed state.

In this context, a striking peculiarity is the redox regulation of the apyrase activity. In oxidized state, the C-terminal R-loop is folded, with a disulfide bond formed between its two cysteines, while it is unfolded in absence of the disulfide bond in the reduced state. When the R-loop is reduced or absent, the T-loop can assume its ATP-protecting conformation, in which the T-loop wraps around the phosphate groups of bound nucleotides, which is presumably the default in SbtB proteins that are lacking the R-loop extension. We found that the apyrase activity is significantly dampened in this state. However, when the R-loop is oxidized, the folding of the T-loop in the ATP-protecting conformation is prevented, which increases the apyrase activity significantly. Since the redox state in oxygenic phototrophs switches between the dark and light, our results indicate that *Sc*SbtB has increased apyrase activity in the dark, when oxidized, resulting in an accelerated turnover from the ATP-bound to the AMP-bound state of SbtB, and thereby to an accelerated closure of the bicarbonate channel SbtA.

Our results imply that the R-loop is reduced enzymatically by a thioredoxin *in vivo*. A hairpin structure similar to the R-loop was previously found in disulfide oxidoreductases (PD: 3GL5) in close vicinity to the catalytic active CPxC motif, underlying the relevance of such segment for regulatory functions, presumably as a redox switch. Our results show that the major *Synechocystis* thioredoxin, TrxA, is able to interact specifically with the oxidized R-loop of *Sc*SbtB. Remarkably, TrxA also shows a light-to-dark response, similar to SbtB, such that its expression is decreased in the dark (Pérez-Pérez et al. 2009; Selim et al. 2021a). Thus, TrxA could be the main SbtB-reducing enzyme in the day-phase, owing to its cellular abundance (Mallén-Ponce et al. 2021). Interestingly, the *trxA* mutant showed a remarkable impairment of the Calvin–Benson–Bassham (CBB) cycle, underlying its importance for the photosynthetic life style of cyanobacteria (Mallén-Ponce et al. 2021). Several CCB enzymes evolved redox-regulated C-terminal extensions formed by a C(V/I)VxVC motif, which are analogous to the CGPxGC motif of the SbtB R-loop, with the same spacing of the cysteines. Strikingly, this motif in CCB enzymes is known as a redox switch to regulate their activity via similar intramolecular disulfide bonds under successive day/night cycles (Michelet et al. 2013; Tamoi et al. 2015). Moreover, the disulfide bonds of CP12 proteins, small CBB-regulatory proteins, are also known to be redox-regulated via TrxA, and even known to form gene clusters with TrxA in cyanobacteria (Hackenberg et al. 2018; McFarlane et al. 2019). Also, a previous study showed that GlgB, one of SbtB targets, is a potential TrxA target (Lindahl & Florencio 2003), which further indicates the importance of TrxA in controlling the central carbon metabolism.

As the R-loop is only found in a subset of SbtB proteins, it seems to be an optional additional regulatory element that is not present in all cyanobacteria. Indeed, it is not involved in direct physical contacts within the SbtA:SbtB complex, but only in controlling apyrase activity. Thus, the architecture of the SbtA:SbtB complex is expected to be the same in all cyanobacteria, independent of the presence or absence of the R-loop extension. Likewise, it is possibly also not a decisive element for the binding to the other SbtB targets, like the c-di-AMP-dependent interaction with the glycogen branching enzyme GlgB (Selim et al. 2021a). Here, the sensing and signaling of c-di-AMP affects glycogen synthesis, leading to reduced glycogen levels in SbtB and c-di-AMP cyclase deficient mutants. During the day, the cells invest energy to accumulate high intracellular bicarbonate concentrations under C_i_-limiting conditions and to maintain the Na^+^-homeostasis required for HCO_3_ transport, to finally synthesize glycogen molecules as a reservoir for the night phases. In this context, the apyrase activity appears to be irrelevant, as there is no functional difference between the SbtB:ATP and the SbtB:AMP complex for the regulation of GlgB, although it cannot be excluded that the redox state of the R-loop has an influence on the direct SbtB:GlgB interaction. Therefore, it is possible that the R-loop is only used to fine-tune the interaction with a subset of targets, possibly only SbtA.

Taking into account all data that we acquired for SbtB and its interaction partners, we propose a refined model for the control of bicarbonate uptake through the SbtA-SbtB system (Fig. 4). Generally, there is always competition of the different adenine nucleotides for SbtB binding, and an equilibrium is found on the basis of the different affinities of these nucleotides and their relative concentrations in a given situation, e.g. in the light or in darkness. However, only the SbtB:AMP complex is able to bind and occlude the SbtA channels; the binding of any other adenine nucleotide disrupts this occlusion via a modulation of the T-loop. During the day, ATP is by far more abundant than AMP and ADP, whereas the concentration of cAMP depends on the CO_2_ supply to the cells. *In vitro*, *Sc*SbtB has the highest affinity towards cAMP, which competes with the other nucleotides. We propose that during the day, most of SbtB is either in the ATP-protecting ATP-bound state (low CO_2_) or in the cAMP-bound state (high CO_2_). Notably, the major adenylyl cyclase in *Synechocystis* (encoded by *slr1991*) is activated by CO_2_ but not by HCO_3_^-^ (Hammer et al. 2006), therefore cAMP levels increase only in presence of CO_2_ (Selim et al. 2018), and would not be influenced by the intracellular levels of HCO_3_^-^ accumulated by CCM. Another fraction of SbtB is also in the c-di-AMP-bound state, thereby linking C_i_-acquisition with glycogen anabolism via GlgB and other potential targets. All these forms of SbtB have only weak affinity to SbtA, thereby keeping the channel in the open conformation. However, when the cells are exposed to darkness, the ATP levels are dropping, increasingly populating the SbtB:AMP state, which can bind to SbtA and closes the HCO_3_^-^ channels. The transition from the SbtB:ATP to the SbtB:AMP complex is additionally aided by the slow basal apyrase activity, and here, the redox regulation of the R-loop comes into play. Upon exposure to darkness, the R-loop is oxidized and thereby upregulates the basal apyrase activity, to speed up the occlusion of SbtA. The fact that the R-loop is not found in all cyanobacteria points at different regulatory needs for the timing of this process, in the different ecological niches populated by cyanobacteria.

**Fig 4:**
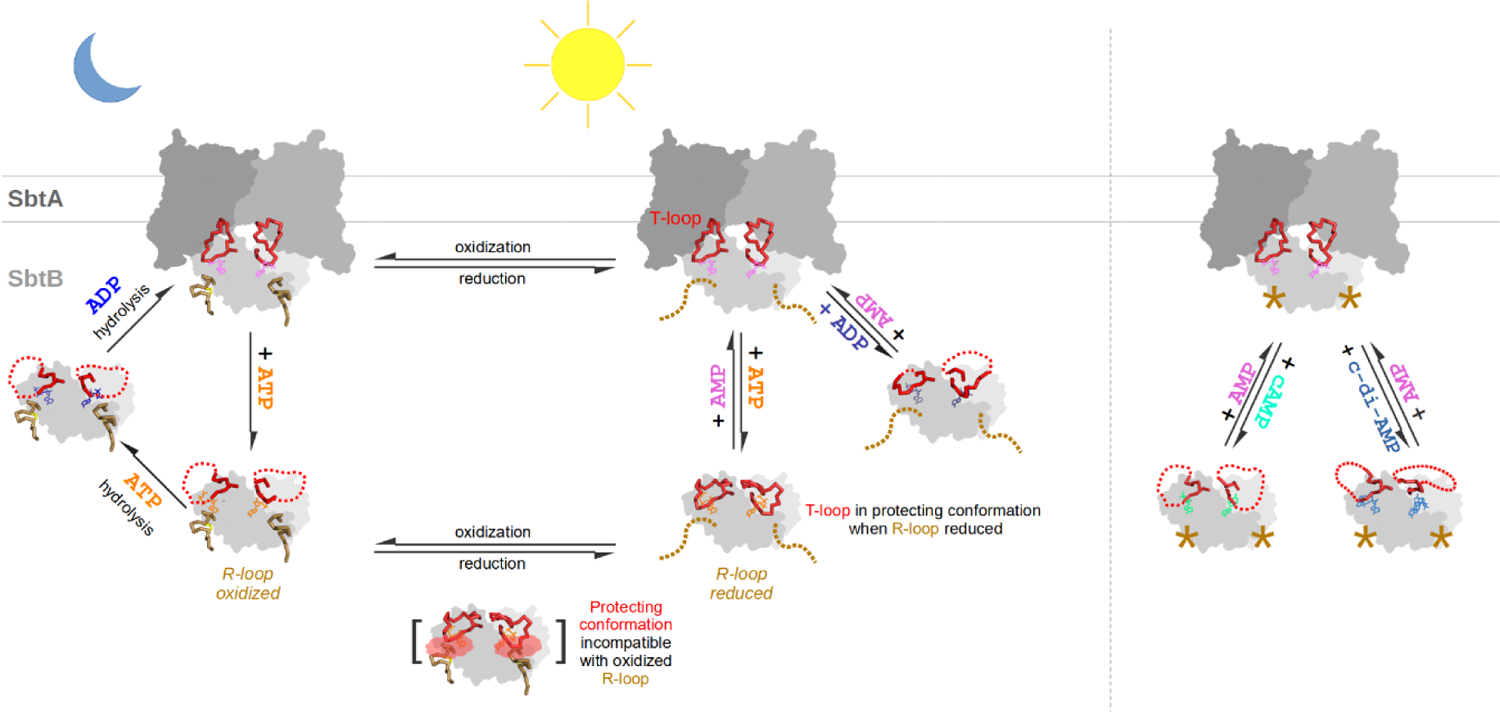
Scheme of nucleotide- and redox-based regulation of the SbtA:SbtB complex. See discussion for details. Asterisks indicate that R-loop state is irrelevant.

## Material and methods

### Generation and purification of recombinant proteins

All the plasmids and primers used in this study are listed in (Table S1). The recombinant C-terminal StrepII-tagged SbtB (wildtype or different variants) proteins from *Synechocystis* sp. PCC 6803 (*Sc*SbtB) or *Nostoc sp.* PCC 7120 (*NsSbtB*) were expressed and purified as previously described (Selim et al. 2018; Lapina et al. 2018) on Strep-Tactin®/Superflow® high-capacity column (IBA), followed by size-exclusion chromatography using the ÄKTA purifier (GE Healthcare) on Superdex^TM 200^ prep-grade column (GE Healthcare). Truncated C-terminal StrepII-tagged *Sc*SbtB protein (SbtB-Δ104), in which the last 6 amino acids (C_105_GPEGC_110_) at the C-terminus is deleted, was constructed using the primer pairs (1256_Fw and 1663_Rv) to generate pASK-IBA3_*Sc*SbtB-Δ104 plasmid. The pASK-IBA3_*Sc*SbtB-Δ104 plasmid was used as a DNA templet to generate the pASK-IBA3_*Sc*SbtB-C105A+C110A and pASK-IBA3_*Sc*SbtB-C105S+C110S plasmids using primer pairs containing the C105+C110 substitution into alanine or serine. For generation of *Sc*SbtB-K40A, *Sc*SbtB-R43A, and *Sc*SbtB-R46A, we used the wildtype pASK-IBA3_*Sc*SbtB-strep plasmid as a DNA templet and primer pairs containing the K40, R43 or R46 substitution into alanine. All of the *Sc*SbtB variants were created using AQUA cloning, in which the linear DNA fragments of different SbtB inserts into the backbone of pASK-IBA3 was transformed into *E. coli* cells (NEB 10-beta). The recombinant N-terminal His_6_-tagged TrxA protein encoded by *slr0623* was expressed and purified as previously described previously for His-tagged proteins (Lapina et al. 2018; Selim et al 2019 and 2020b) on Ni^2+^-NTA column.

### Phosphate release Assay

After an overnight dialysis in phosphate free reaction buffer [50 mM Tris/HCl, 200 mM NaCl; pH 7.4] with or without 1 mM MgCl_2_ at 4°C, 50 or 100 µM of respective SbtB (wildtype or its variants) preparation (monomeric concentration) was mixed with either 100-300 µM ATP or ADP in a 1.5 mL reaction tube. Samples were incubated at 28°C in a rotary shaker [Thriller®, peqlab] for overnight or as indicated. After the incubation time, 250 µL samples were taken (either before (T_0_) or after the end of incubation (T_Final_)), proteins were separated from each sample by centrifugation through Amicon ® Ultra Centrifugal Filters (Merck, Millipore Ltd.) at 21.000 x g for 10 min at 4°C. 20 µL of the flow-through were diluted in 80 µL of phosphate free reaction buffer and used to determine phosphate concentration using the ab65622 Phosphate Assay Kit (Colorimetric) [abcam^®^] according to manufacturer’s instructions. To exclude the influence of spontaneous hydrolysis of ATP and ADP under our tested conditions, negative controls for the nucleotides without the proteins in the same phosphate free buffer and incubated for the same periods were used. The negative controls revealed a marginal phosphate release, which was further subtracted from the released phosphate at T_Final_, if there was any. Released phosphate was calculated by subtracting T_0_ values and the spontaneous hydrolysis for the nucleotides from T_Final_ values. For reproducibility, all measurements were performed at least in triplicates with different purification batches of recombinant proteins as indicated. To test the influence of the metal ions on the apyrase activity of SbtB, 5 mM of respective tested metals was used instead of MgCl_2_ and compared to metal free buffer.

### Crystallization, Crystal handling, Data collection, and Structure determination

All crystallization experiments were performed at 20 °C using the vapor diffusion method in 96-well sitting-drop plates, with crystallization drops containing 300 nl reservoir solution and 300 nl protein solution equilibrating against 50 μl reservoir solution. Apo crystals of wildtype *Sc*SbtB were grown as described before (Selim et al. 2018), with a protein solution containing 15–20 mg/mL of *Sc*SbtB, 50 mM Tris/HCl (pH 8.0), 300 mM NaCl, and 0.5 mM EDTA, and a reservoir solution containing 0.2 M sodium acetate, 0.1 M HEPES pH 7.5, and 20% (w/v) PEG 3000.

For co-crystallization attempts of wildtype *Sc*SbtB protein with ADP or ATP, 3-5 mM of ADP or ATP were added to the protein solution and full crystallization screens performed as described (Selim et al. 2018). For co-crystallization of SbtB^defR^ variants with ADP (*Sc*SbtB Δ104 variant) and ATP (*Sc*SbtB C105A+C110A variant), 2 mM of ADP or ATP were added to the protein solution and full crystallization screens were performed as described (Selim et al. 2018). SbtB^defR^:ADP co-crystals (*Sc*SbtB Δ104 variant) were obtained in the Molecular Dimensions Morpheus screen condition B9, which is composed of 0.09 M halogen (stock solution: 0.3 M sodium fluoride; 0.3 M sodium bromide; 0.3 M sodium iodide), 0.1 M buffer system 3 pH 8.5 (Tris [base]; Bicine), and 50% (v/v) precipitant mixture 1 (stock solution: 40% (v/v) PEG500; MME; 20% (w/v) PEG 20000). SbtB^defR^:ATP co-crystals (*Sc*SbtB C105A+C110A variant) were obtained with a reservoir solution composed of 0.1 M MES pH 6.5 and 25% (w/v) PEG 1000.

ADP and ATP soaking experiments were performed as described previously for c-di-AMP (Selim et al. 2021a). Briefly, trigonal apo-*Sc*SbtB crystals were soaked in a droplet of reservoir solution supplemented with 2 mM of either ADP or ATP for indicated time spans prior to cryo protection. SbtB^defR^:ADP crystals were loop-mounted directly from the crystallization plates; all other crystals were briefly transferred to a droplet of reservoir solution supplemented with either 20% PEG 400 or glycerol for cryo protection. All crystals were flash-frozen in liquid nitrogen and diffraction data were collected with a wavelength of 1 Å at 100 K on a PILATUS 6M-F detector at beamline X10SA of the Swiss Light Source (PSI, Villigen, Switzerland). All data were indexed, integrated, and scaled using XDS (Kabsch 2010).

All structures were solved based on the trigonal apo-*Sc*SbtB structure (PDB: 5O3P) using either difference Fourier methods (for the structures in space group P3_2_) or molecular replacement via MOLREP (Vagin & Teplyakov 2010) (for structures in P4_1_). After initial rigid body refinement with REFMAC5 (Murshudov et al. 2011), the structures were rebuilt and completed by cyclic manual modeling with Coot (Emsley et al. 2010) and refinement with REFMAC5. Crystals of wildtype *Sc*SbtB obtained in co-crystallization attempts with ADP and ATP revealed to have AMP bound in all binding sites and were not regarded further. Data collection and refinement statistics are shown in (Table S2). Structural representations were prepared using PyMol.

### BACTH assay

All the plasmids and primers used for BACTH assay are listed in (Table S1). Plasmid construction, cell cultivation, and experimental procedure of BACTH assay were performed as described previously (Selim et al. 2021a) on X-Gal and MacConkey agar plates supplemented with ampicillin (100 μg/ml), kanamycin (50 μg/ml), and IPTG (1 mM). The T25-SbtB fusion with an empty pUT18 vector was used as negative control, while the leucine zipper interaction was used as positive control. As described previously, GlgB and GlgC were used as positive and negative controls for SbtB interaction, respectively (Selim et al. 2021a). The *E. coli* BTH101 (Euromedex) was used for BACTH assays. The BACTH assays were performed at least two-times with three independent *E. coli* colonies to confirm the reproducibility and the specificity of the SbtB and its variant to interact with TrxA.

### Pulldowns and mass spectrometry analysis

For pulldown experiments, 10 µM of either strep-tagged SbtB or His-tagged TrxA protein was pre-incubated at 28°C for 15 min on 15 µl of either magnetic MagStrep “type3” XT beads (IBA GmbH, Göttingen) or Ni^2+^-NTA magnetic Beads (MagBeads; Genaxxon), respectively. Next, 10 µM of the interacting partner was added to the immobilized protein and allowed to interact for another 15 min at 28°C. To get rid of the excess unbound proteins, the magnetic beads were extensively washed for at least four times using either 200 µl of Strep Washing Buffer (100 mM Tris/HCl, 200 mM NaCl, 1 mM EDTA; pH 8.0) Ni-NTA Washing Buffer (50 mM Na_2_HPO_4_, 200 mM NaCl, 10 mM imidazole; pH 8.0) for Strep-tagged immobilized SbtB and His-tagged immobilized TrxA, respectively. 20 µl of the fourth washing steps were used for SDS-gel application to make sure that the unbound proteins were successfully washed away. Finally, the elution was performed by the addition of 20 µl of either Strep-BXT Biotin Elution Buffer (IBA) or His-elution Buffer (50 mM Na_2_HPO_4_, 200 mM NaCl, 500 mM imidazole; pH 8.0). As negative control empty beads were incubated with 10 µM of recombinant TrxA-His over magnetic MagStrep beads or 10 µM of recombinant Streptagged SbtB over Ni^2+^-NTA magnetic beads to further exclude the unspecific binding of the proteins to the magnetic beads. For analysis of interacting partners of pulldown experiments, 20 µl of each fraction were applied to a 12% SDS-PAGE gel. For detection of the protein, which coeluted with the immobilized protein, we used western blotting analysis using anti-Strep-tag II antibody (abcam) for the detection of recombinant Strep-tagged SbtB, which coeluted with His-TrxA or monoclonal anti-polyHistidine-Peroxidase antibody (Sigma-Aldrich) for the detection of recombinant His-tagged TrxA, which coeluted with Strep-SbtB.

For mass spectrometry analysis to identify if SbtB contains a protein contamination that possesses ATPase activity, recombinant purified SbtB protein from *E. coli* cells was in gel digested with trypsin, then LC-MS/MS analysis was performed on a Proxeon Easy-nLC coupled to QExactive HF using 60 min gradient. Processing of data was achieved using MaxQuant software (Version 1.6.7.0. with integrated Andromeda Peptid search engine). The spectra were searched against *E. coli* database (UP000000625_83333_complete_2019-12-11), *Synechocystis* database (UP000001425_1111708_complete_2019-02-13) and sequences for SbtB.

## Acknowledgments

This work was supported by the German research foundation (DFG) within the priority program SPP1879 to K.F., the Cluster of Excellence (EXC 2124) “Controlling Microbes to Fight Infections” to K.A.S., and by institutional funds of the Max Planck Society. We are grateful to N. Neumann and H. Grenzendorf (IMIT, Tübingen University) for the excellent assistance, the staff of beamline X10SA/SLS, the Proteome Center (Tübingen University), and L. Lo-Presti for critical scientific and linguistic editing of the manuscript. Furthermore, we would like to acknowledge the infrastructural support by the Cluster of Excellence “Controlling Microbes to Fight Infections” (EXC 2124–390838134) of the DFG.

## Author contributions

K.A.S. and K.F. conceived and initiated the project. K.A.S. and M.D.H. designed the experiments. K.A.S. and M.H. performed the biochemical experiments. R.A. performed crystallographic sample preparation and diffraction data collection, K.A.S. solved the crystal structures, and M.D.H. supervised the structural analysis. K.A.S., H.Z. and M.D.H evaluated and interpreted the results. K.A.S., K.F. and M.D.H wrote the manuscript. All authors approved the final version of the manuscript.

## Data and materials availability

Crystallography, atomic coordinates, and structure factors have been deposited in the Protein Data Bank, www.wwpdb.org (PDB ID codes: 7R2Y, 7R2Z, 7R30, 7R31, and 7R32).

## Supplementary Materials

**Fig. S1.**
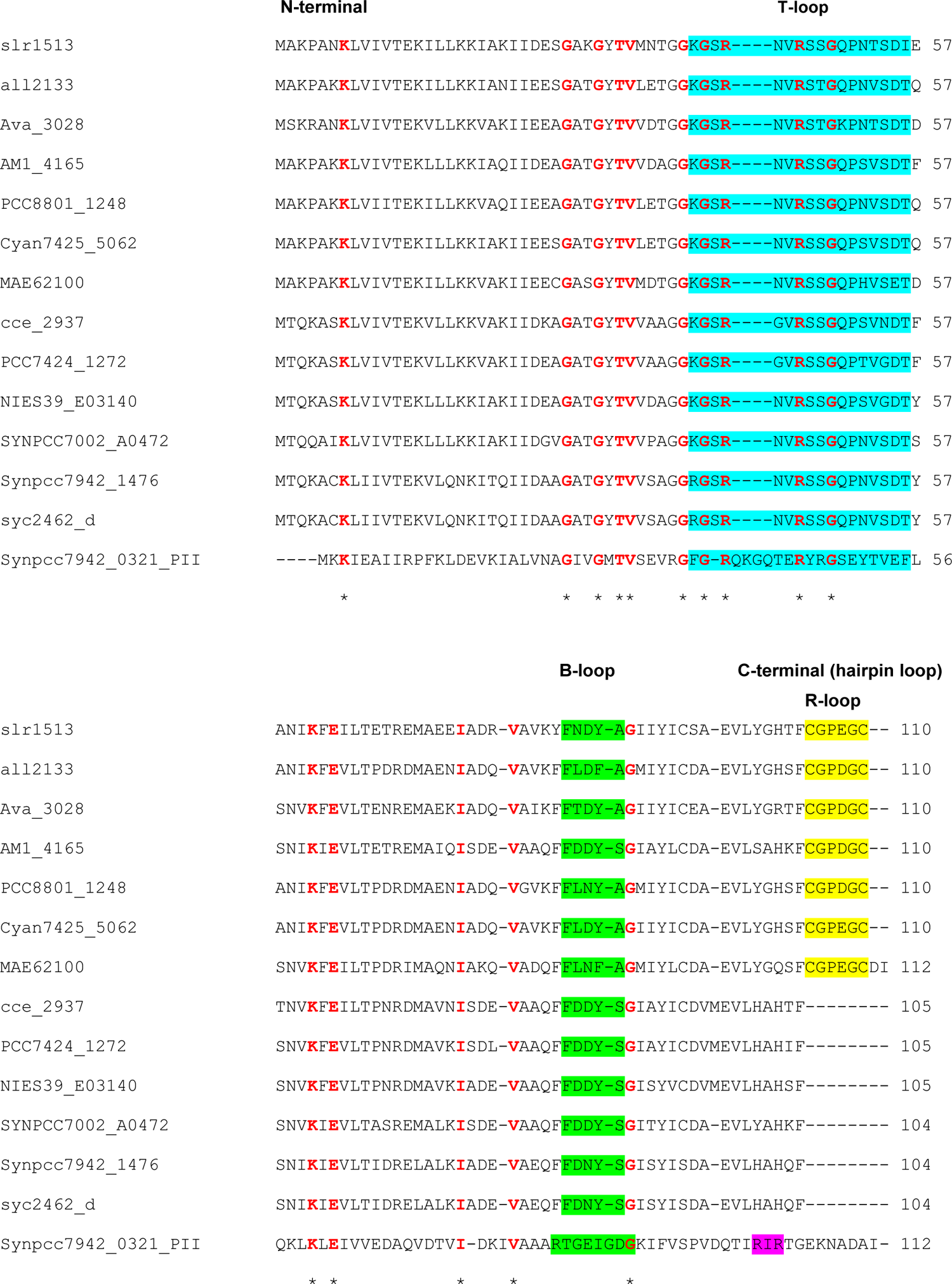
Multiple sequence alignment of SbtB homologs in cyanobacteria. SbtB sequences were extracted from CyanBase via a BLAST search using the protein sequences of *Sc*SbtB and *Ns*SbtB (encoded by *slr1513* and *all2133*, respectively) as query and aligned with canonical PII protein (encoded by *Synpcc7942_0321*_PII) using Clustal Omega. Residues highly conserved in canonical PII proteins are in red and indicated with asterisks. The T-loop and B-loop of SbtB and PII proteins are highlighted in blue and green, respectively. In the C-terminal region, the PII arginine fingerprint motif (RxR), which is known to coordinate the β- and γ-phosphates of ATP or ADP, is highlighted in pink. The C-terminal hairpin loop in SbtB proteins, which forms a disulfide bond between Cys105 and Cys110 and therefore we termed R-loop (standing for redox-regulated loop), is highlighted in yellow.

**Fig. S2.**
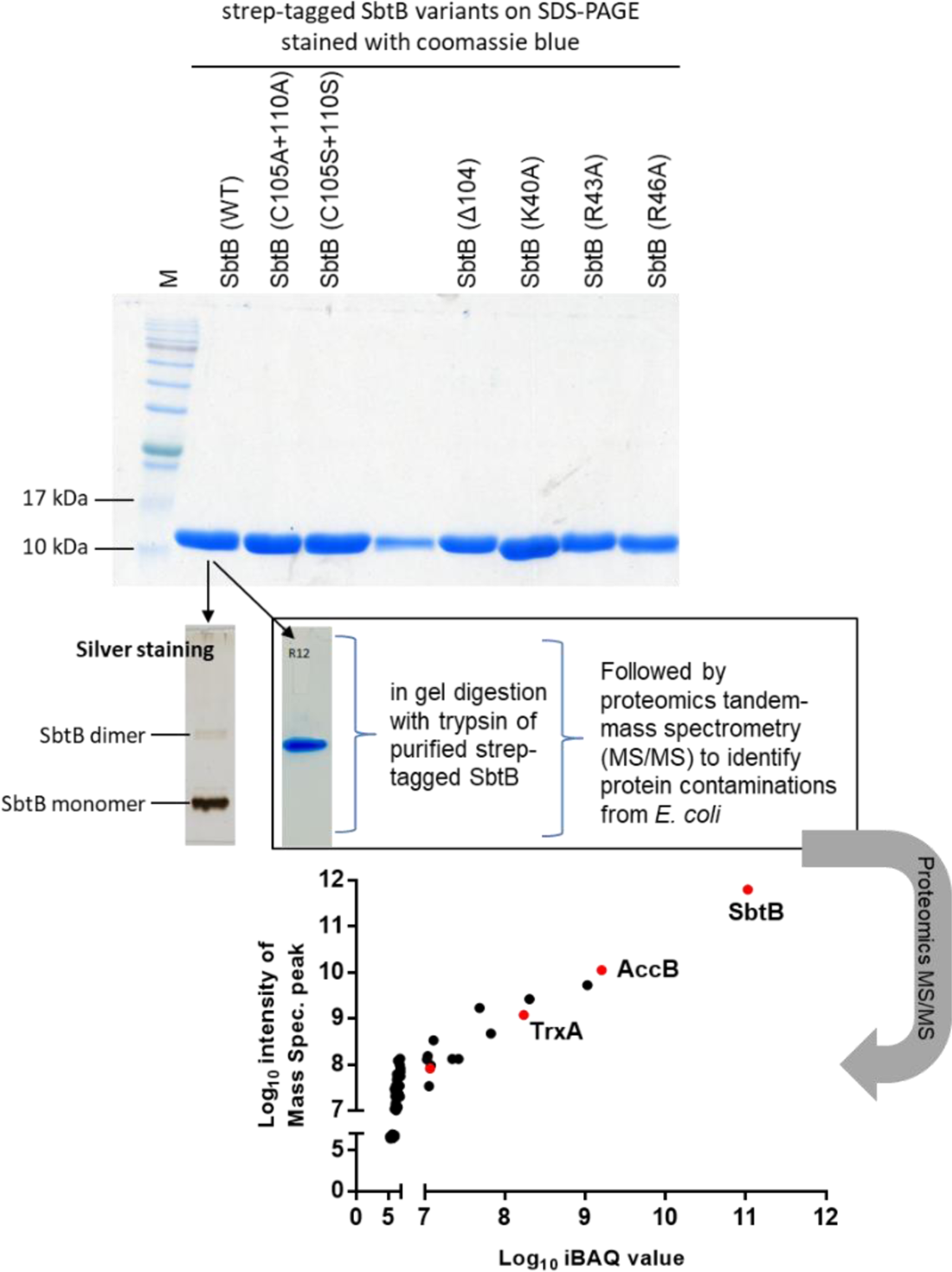
SDS-PAGE stained by Coomassie blue for strep-tagged SbtB variants used in this study after purification from *E. coli* expressing the respective protein. SDS-PAGE showed high degree of purity for all purified SbtB variants. To check for residual protein contaminations from *E. coli*, the wildtype SbtB (WT) was further checked by silver staining, which is more sensitive than Coomassie blue stain, and moreover it was subjected to tandem-mass spectrometry (MS/MS) to identify the *E. coli* proteins, which coeluted with wildtype SbtB (check proteomic dataset associated with this manuscript). The identified proteins were sorted based on iBAQ values of significantly enriched proteins and plotted against the intensity of MS peaks of the identified/defined peptides. The red dots refer to SbtB, thioredoxin-1 (TrxA), glutaredoxin-4 (GrxD), and biotin carboxyl carrier protein of acetyl-CoA carboxylase (AccB). Biotinylated proteins (AccB) are common contaminant of strep-tag purifications. TrxA and GrxD are of special interest as potential targets of SbtB to break the R-loop (check the main text).

**Fig. S3.**
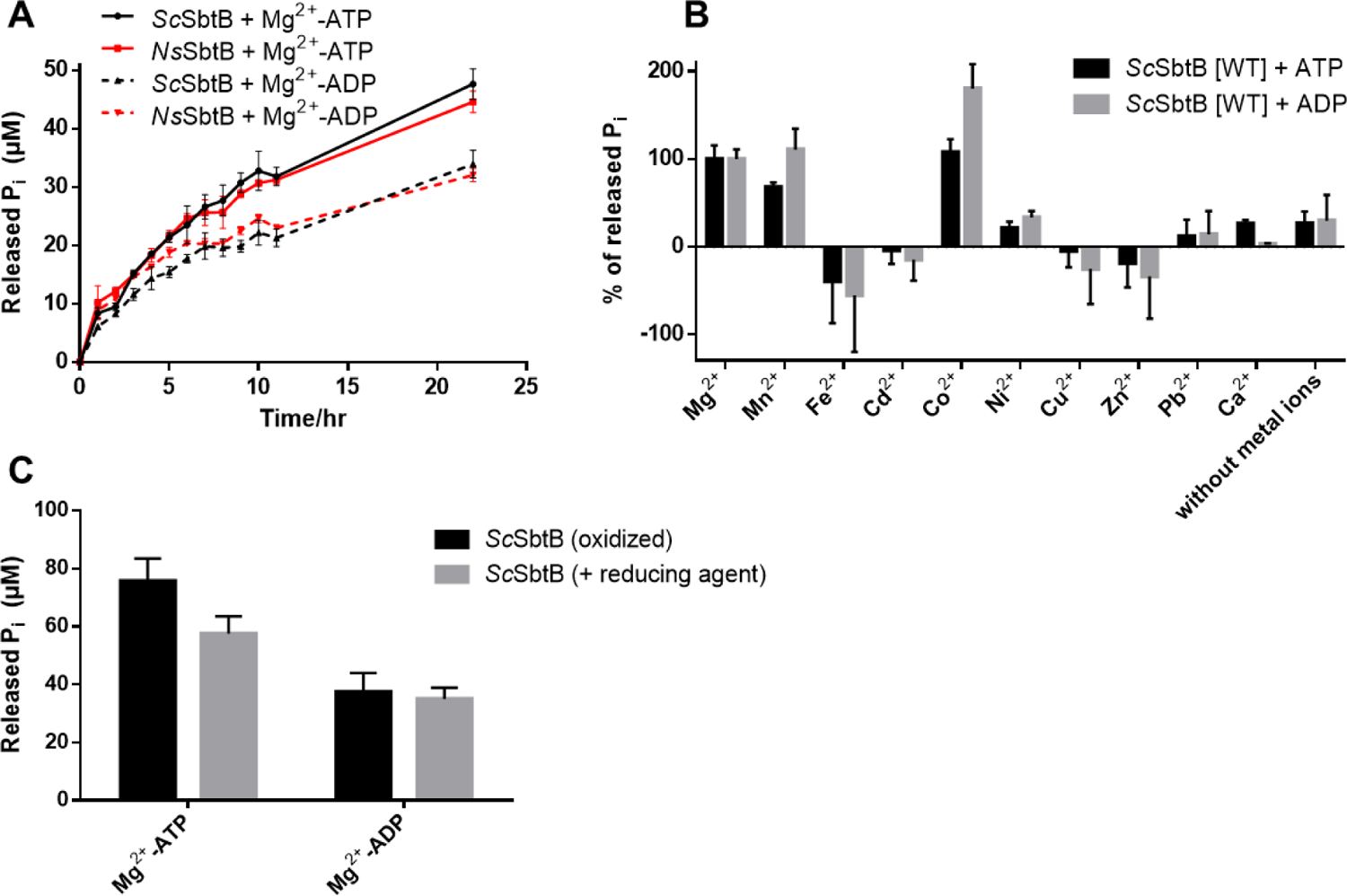
Apyrase activity of SbtB proteins via phosphate release assay. (A) Time course for slow ATP and ADP hydrolysis via *Sc*SbtB and *Ns*SbtB, revealing that ATP and ADP hydrolysis are a common trait among SbtB proteins. The released inorganic phosphate (P_i_) is shown in µM. (B) Metal influence on *Sc*SbtB apyrase activity, relative to wildtype *Sc*SbtB-activity in presence of Mg^2+^ (100%). The assay was performed in presence of 5 mM of the respective metals. Negative values are indicative of heavily protein precipitation. The assay indicated that Mn^2+^, Mg^2+^ and Co^2+^ could be used as metal ions by *Sc*SbtB. The only metal which can be found in excess inside cells is Mg^2+^, and since high Co^2+^ concentrations is not of physiological relevance, therefore we concluded that Mg^2+^ is most likely the metal used by *Sc*SbtB. (C) Influence of reducing agent on *Sc*SbtB apyrase activity compared to *Sc*SbtB under oxidizing conditions, showing that addition of 1 mM TECP does not influence on *Sc*SbtB apyrase activity.

**Fig. S4.**
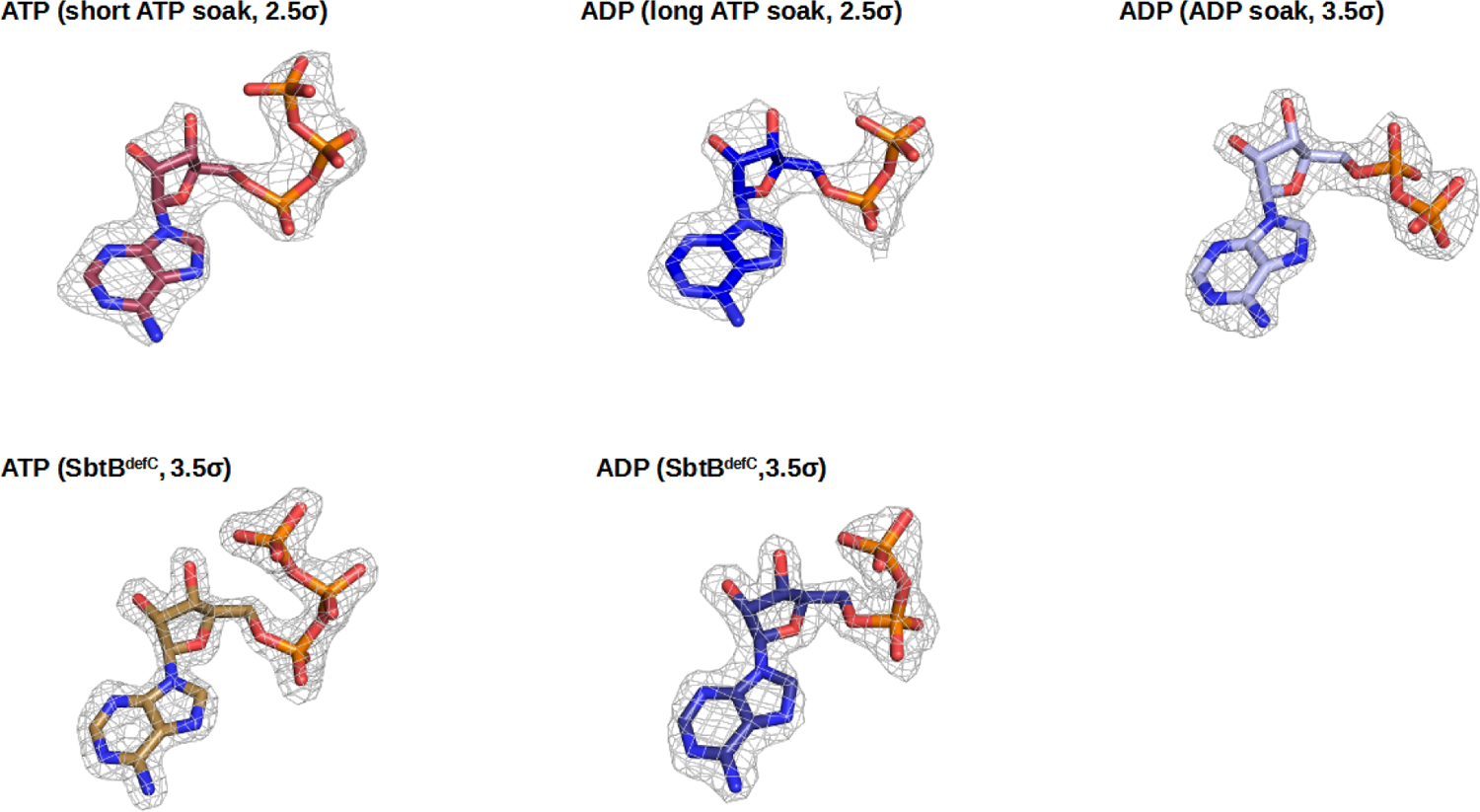
Electron densities of nucleotides.

**Fig. S5.**
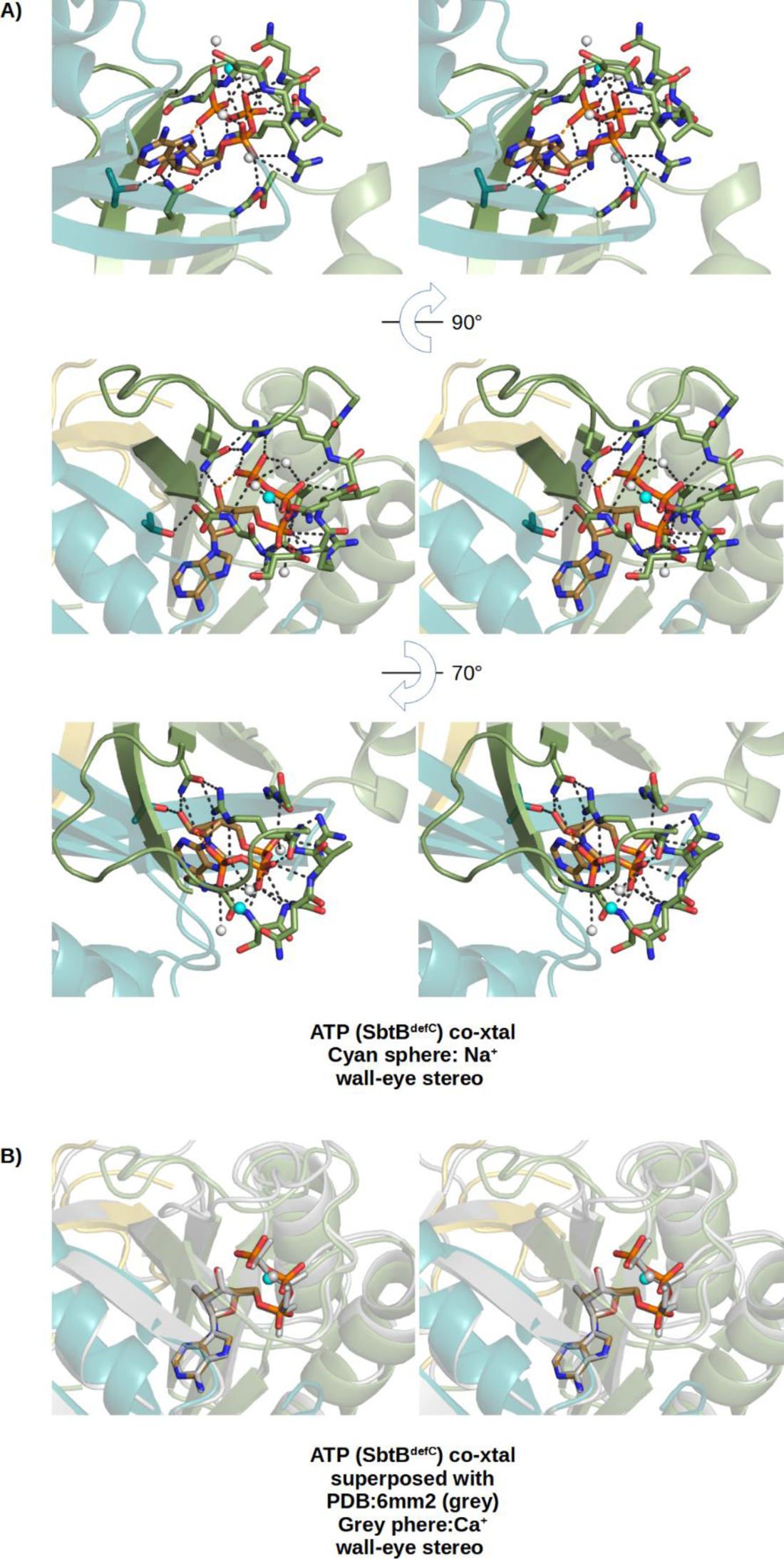
Stereo views of the SbtB^defR^:ATP complex. A) The ATP binding mode is shown in the same orientation as in (Fig. 3), plus two additional orientations, in stereo. B) Stereo superposition of the SbtB^defR^:ATP complex to the SbtB:ATP complex from *Cyanobium* sp. PCC7001 (PDB: 6MM2).

**Fig. S6.**
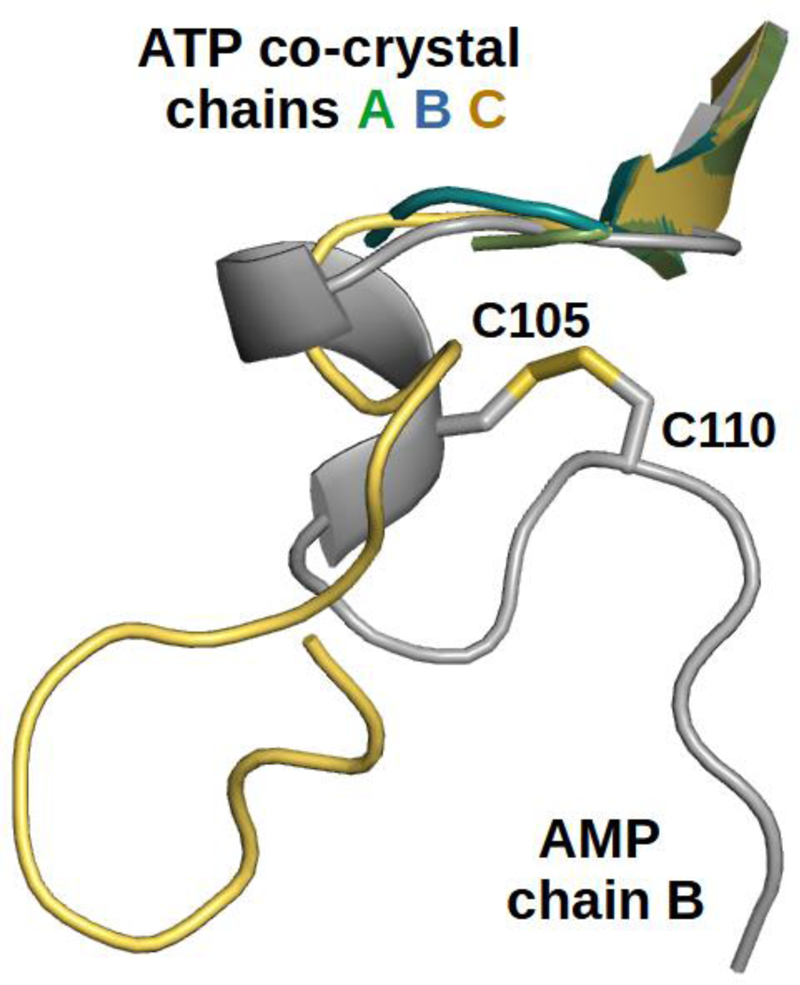
Comparison of folded and unfolded R-loop. The R-loops of the three chains of the SbtB^defR^:ATP co-crystal structure, in which the two R-loop cysteines were substituted by alanine to mimick the reduced state, are superimposed to the oxidized R-loop in the SbtB:AMP co-crystal structure. Obviously, the fold of the oxidized state is not assumed without the disulfide bond and the R-loop completely disordered in two of the three chains.

**Fig. S7.**
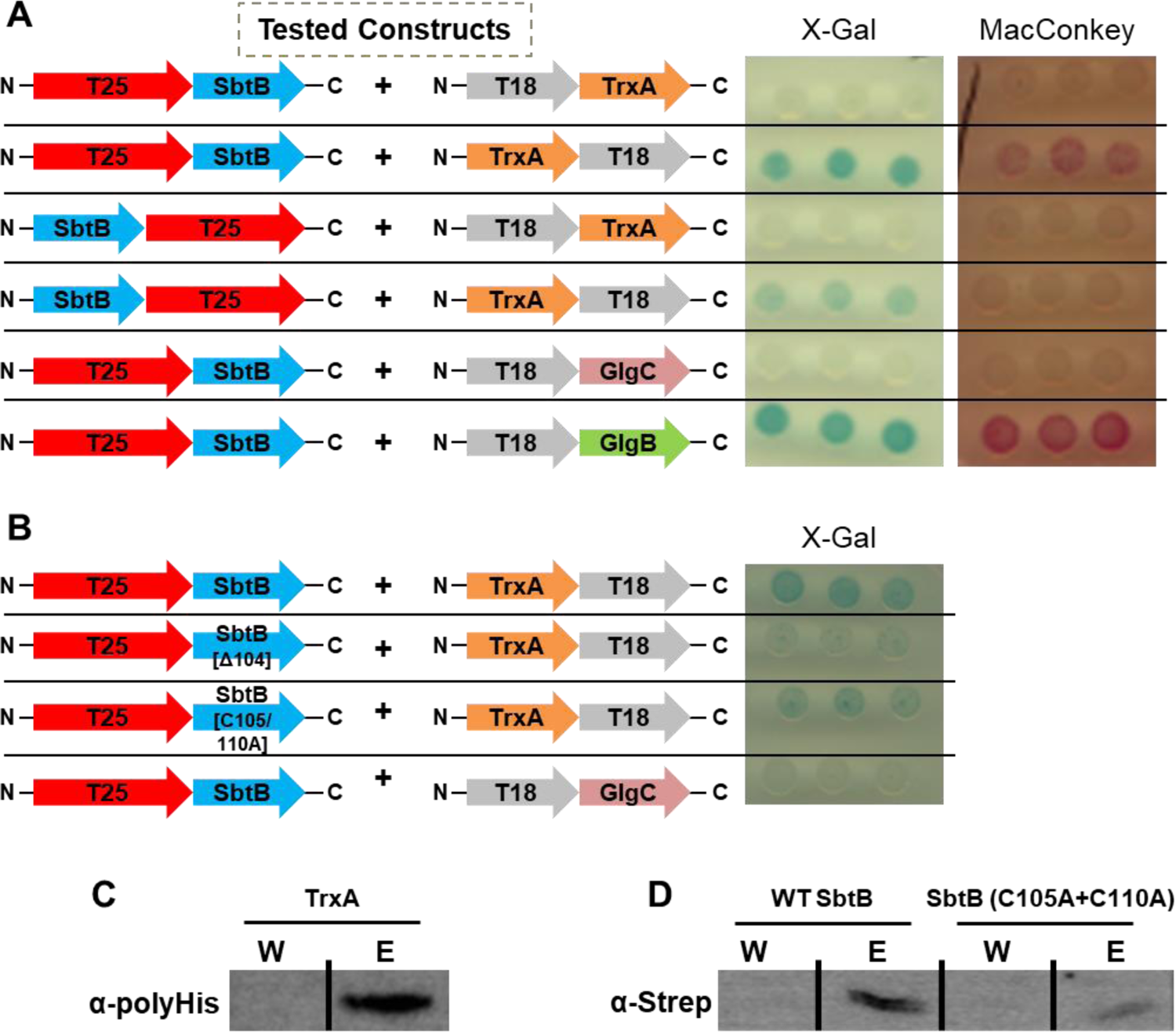
Analysis of the interaction between SbtB and TrxA via bacterial two hybrid assay (BACTH) and pulldown assays. (A) The BACTH assay was performed using *E. coli* cells expressing either N- or C-terminal fusion of Cya-T25 subunit to SbtB together with either N- or C-terminal fusion of Cya-T18 subunit to TrxA, as indicated, on X-Gal or MacConkey reporter plats. N-terminal fusion of Cya-T25 subunit to SbtB together with N-terminal fusion of Cya-T18 subunit with either GlgB or GlgC, was used as positive and negative control, respectively (13). (B) Influence of mutating SbtB R-loop residues on TrxA interaction. The BACTH assay was performed using *E. coli* cells expressing N-terminal fusion of Cya-T18 subunit with TrxA together with N-terminal fusion of Cya-T28 subunit of either wildtype SbtB, or SbtB-(Δ104), or SbtB-(C105A+C110A) as indicated, on X-Gal reporter plat. Positive interaction is evidenced by appearance of a blue or red color on X-Gal or MacConkey reporter plates, respectively. The assay was done using 3-independent/freshly transformed *E. coli* cells, for at least three times to ensure reproducibility. (C and D) Immunoblot blot analysis of SbtB and TrxA interaction in last wash (W) and elution (M) fractions. (C) Strep-tagged SbtB was immobilized and the coelution of TrxA was checked using α-polyHis antibody. (D) His-tagged TrxA was immobilized on Ni^2+^-NTA and the coelution of wildtype SbtB or its variant (C105A+C110A) was checked using α-strep antibody.

---

**Table S1.**
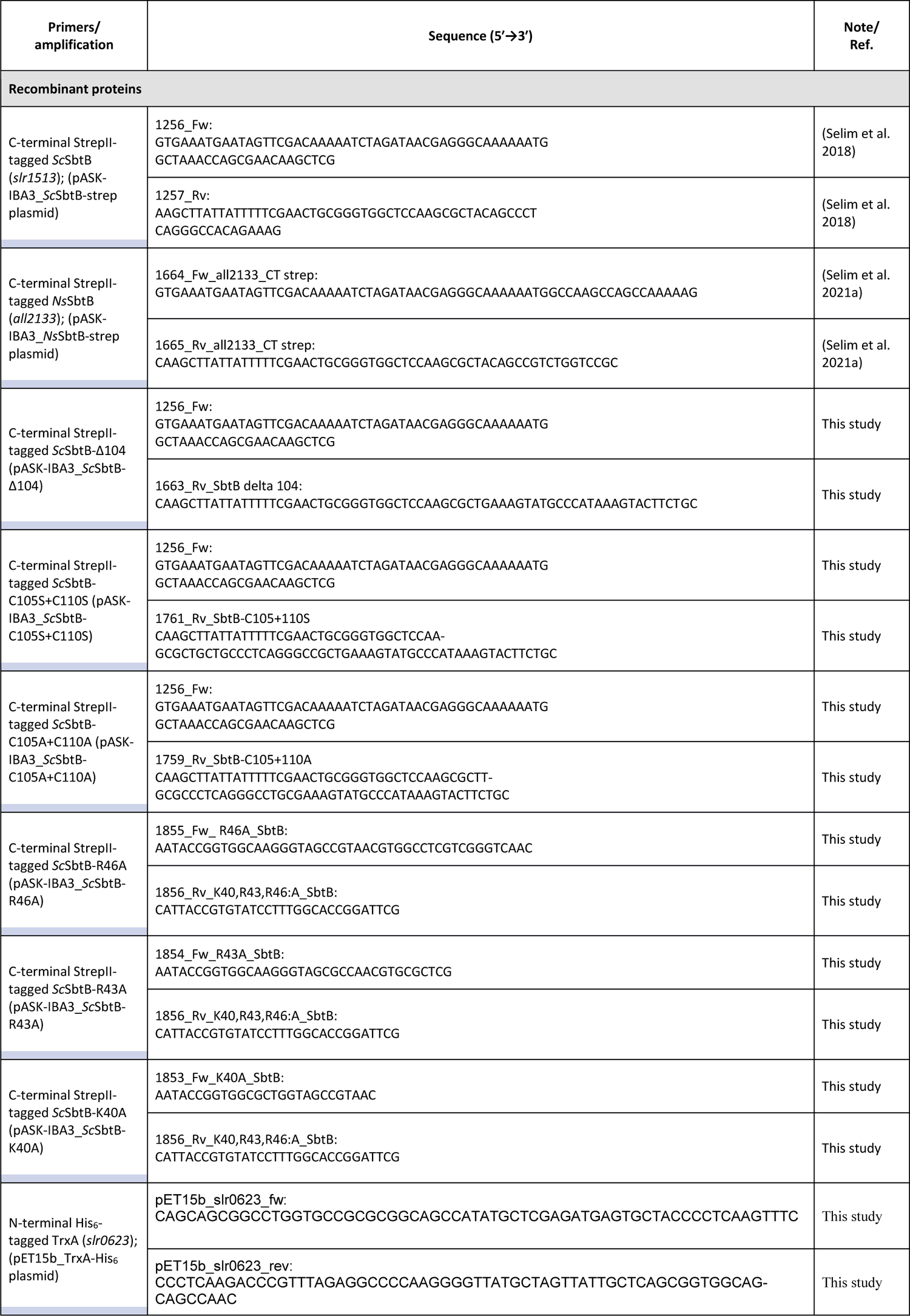
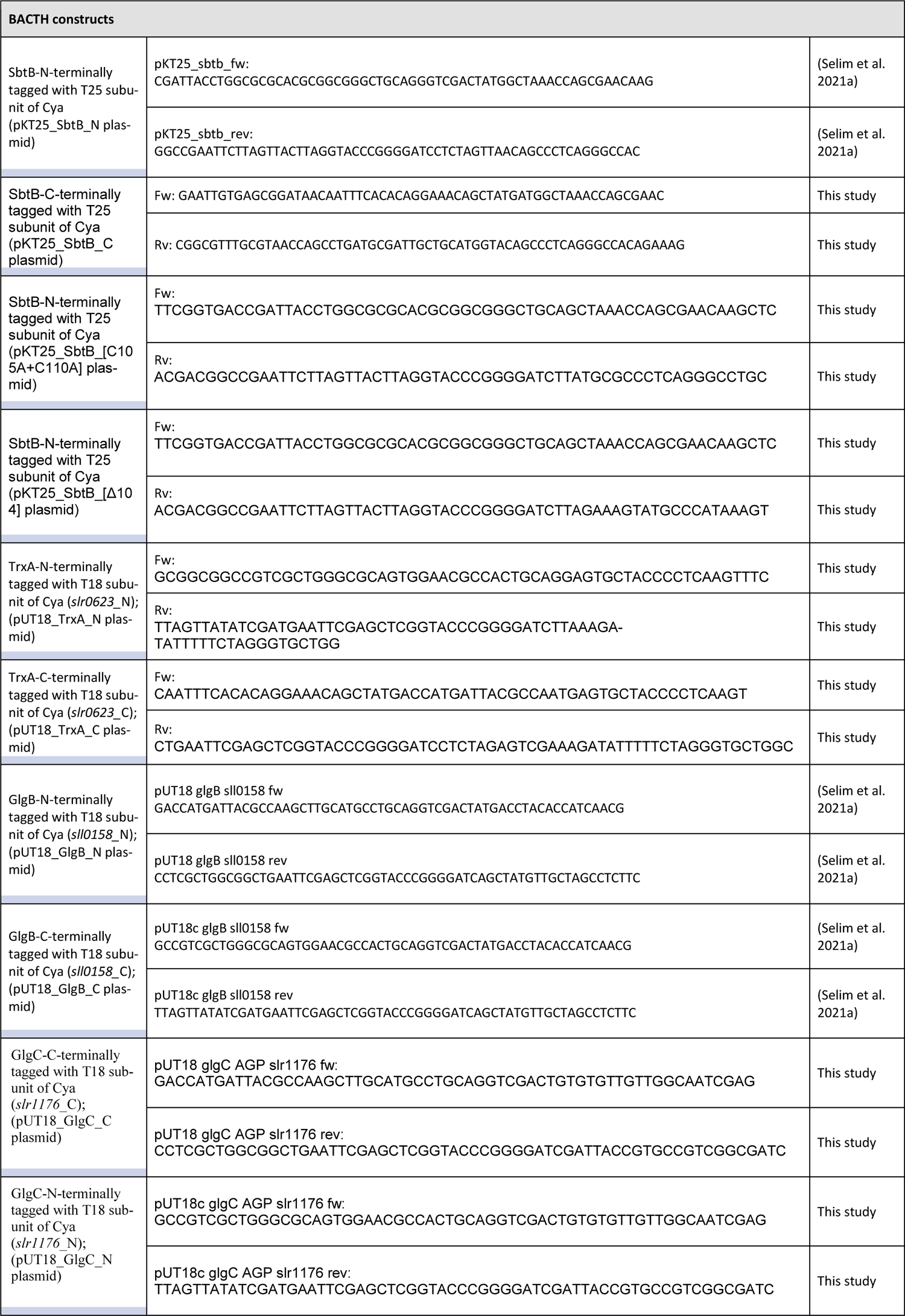
**Primers and Plasmids**

**Table S2.**
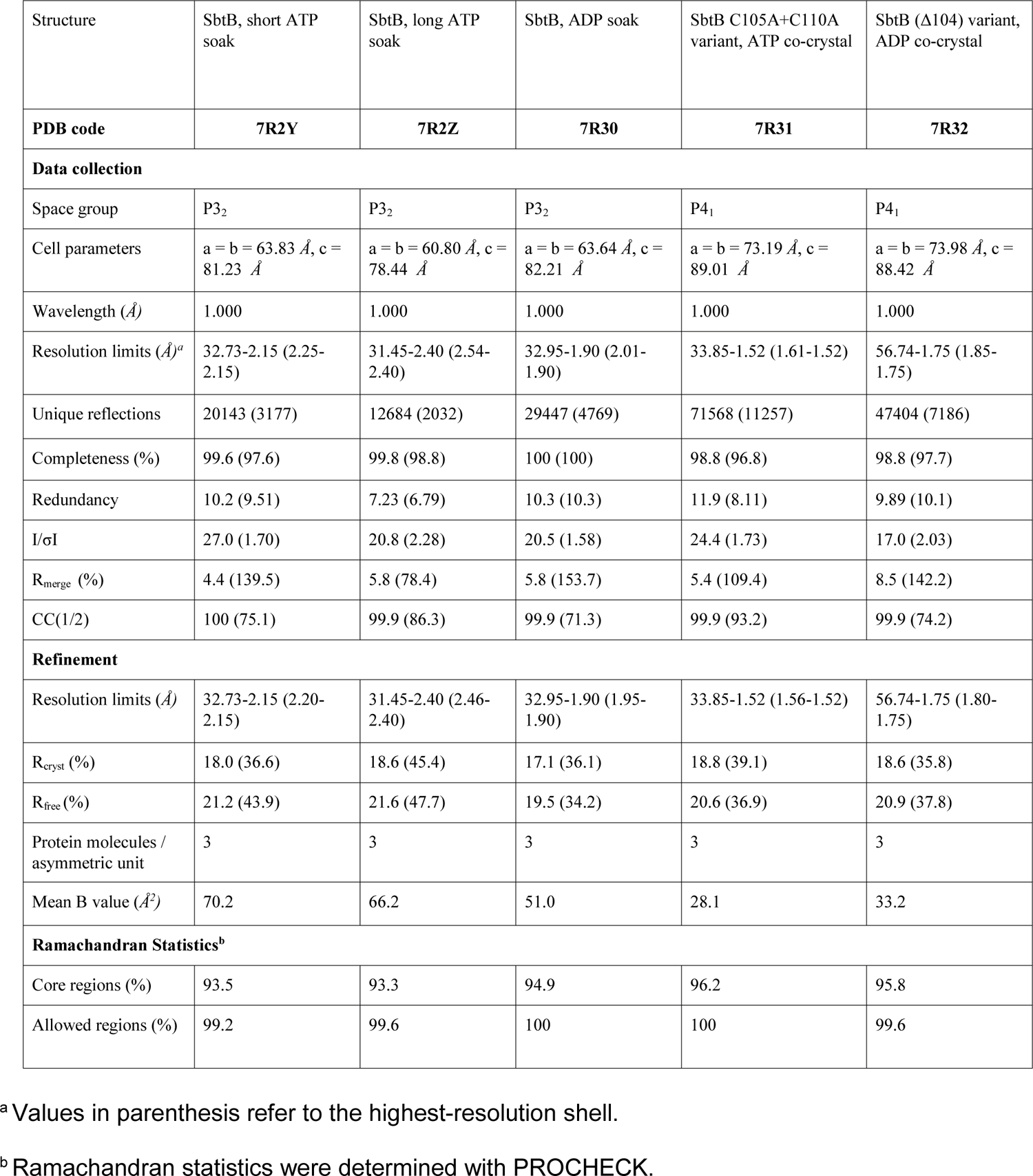
Data Collection and Refinement Statistics

