## Supplementary Information for "Carbon signaling protein SbtB possesses redox-regulated apyrase activity to facilitate regulation of bicarbonate transporter SbtA"

**This PDF file includes:**

Figs. S1 to S7
Tables. S1 to S2

**N-terminal** **T-loop**

```
slr1513               MAKPANKLVIVTEKILLKKIAKIIDESGAKGYTVMNTGGKGSR----NVRSSGQPNTSDIE 57
all2133               MAKPAKKLVIVTEKILLKKIANIIEESGATGYTVLETGGKGSR----NVRSTGQPNVSDTQ 57
Ava_3028              MSKRANKLVIVTEKVLLKKVAKIIEEAGATGYTVVDTGGKGSR----NVRSTGKPNTSDTD 57
AM1_4165              MAKPAKKLVIVTEKLLLKKIAQIIDEAGATGYTVVDAGGKGSR----NVRSSGQPSVSDTF 57
PCC8801_1248          MAKPAKKLVIITEKILLKKVAQIIEEAGATGYTVLETGGKGSR----NVRSSGQPNVSDTQ 57
Cyan7425_5062         MAKPAKKLVIVTEKILLKKIAKIIEESGATGYTVLETGGKGSR----NVRSSGQPSVSDTQ 57
MAE62100              MAKPAKKLVIVTEKILLKKVAKIIEECGASGYTVMDTGGKGSR----NVRSSGQPHVSETD 57
cce_2937              MTQKASKLVIVTEKVLLKKVAKIIDKAGATGYTVVAAGGKGSR----GVRSSGQPSVNDTF 57
PCC7424_1272          MTQKASKLVIVTEKVLLKKVAKIIDEAGATGYTVVAAGGKGSR----GVRSSGQPTVGDTF 57
NIES39_E03140         MTQKASKLVIVTEKLLLKKIAKIIDEAGATGYTVVDAGGKGSR----NVRSSGQPSVGDTY 57
SYNPCC7002_A0472      MTQQAIKLVIVTEKLLLKKIAKIIDGVGATGYTVVPAGGKGSR----NVRSSGQPNVSDTS 57
Synpcc7942_1476       MTQKACKLIIVTEKVLQNKITQIIDAAGATGYTVVSAGGRGSR----NVRSSGQPNVSDTY 57
syc2462_d             MTQKACKLIIVTEKVLQNKITQIIDAAGATGYTVVSAGGRGSR----NVRSSGQPNVSDTY 57
Synpcc7942_0321_PII   ----MKKIEAIIRPFKLDEVKIALVNAGIVGMTVSEVRGFG-RQKGQTERYRGSEYTVEFL 56
                            *                    *  * **    * * *      *  *
```

**B-loop** **C-terminal (hairpin loop)**

**R-loop**

```
slr1513               ANIKFEILTETREMAEEIADR-VAVKYFNDY-AGIIYICSA-EVLYGHTFCGPEGC-- 110
all2133               ANIKFEVLTPDRDMAENIADQ-VAVKFFLDF-AGMIYICDA-EVLYGHSFCGPDGC-- 110
Ava_3028              SNVKFEVLTENREMAEKIADQ-VAIKFFTDY-AGIIYICEA-EVLYGRTFCGPDGC-- 110
AM1_4165              SNIKIEVLTETREMAIQISDE-VAAQFFDDY-SGIAYLCDA-EVLSAHKFCGPDGC-- 110
PCC8801_1248          ANIKFEVLTPDRDMAENIADQ-VGVKFFLNY-AGMIYICDA-EVLYGHSFCGPDGC-- 110
Cyan7425_5062         ANIKFEVLTPDRDMAENIADQ-VAVKFFLDY-AGIIYICDA-EVLYGHSFCGPEGC-- 110
MAE62100              SNVKFEILTPDRIMAQNIAKQ-VADQFFLNF-AGMIYLCDA-EVLYGQSFCGPEGCDI 112
cce_2937              TNVKFEILTPNRDMAVNISDE-VAAQFFDDY-SGIAYICDVMEVLHAHTF-------- 105
PCC7424_1272          SNVKFEVLTPNRDMAVKISDL-VAAQFFDDY-SGIAYICDVMEVLHAHIF-------- 105
NIES39_E03140         SNVKFEVLTPNRDMAVKIADE-VAAQFFDDY-SGISYVCDVMEVLHAHSF-------- 105
SYNPCC7002_A0472      SNVKIEVLTASREMALKISDE-VAAQFFDDY-SGITYICDA-EVLYAHKF-------- 104
Synpcc7942_1476       SNIKIEVLTIDRELALKIADE-VAEQFFDNY-SGISYISDA-EVLHAHQF-------- 104
syc2462_d             SNIKIEVLTIDRELALKIADE-VAEQFFDNY-SGISYISDA-EVLHAHQF-------- 104
Synpcc7942_0321_PII   QKLKLEIVVEDAQVDTVI-DKIVAAARTGEIGDGKIFVSPVDQTIRIRTGEKNADAI- 112
                         * *           *    *          *
```

**Fig. S1. Multiple sequence alignment of SbtB homologs in cyanobacteria.** SbtB sequences were extracted from CyanBase via a BLAST search using the protein sequences of *Sc*SbtB and *Ns*SbtB (encoded by *slr1513* and *all2133*, respectively) as query and aligned with canonical PII protein (encoded by *Synpcc7942_0321*_PII) using Clustal Omega. Residues highly conserved in canonical PII proteins are in red and indicated with asterisks. The T-loop and B-loop of SbtB and PII proteins are highlighted in blue and green, respectively. In the C-terminal region, the PII arginine fingerprint motif (RxR), which is known to coordinate the β- and γ-phosphates of ATP or ADP, is highlighted in pink. The C-terminal hairpin loop in SbtB proteins, which forms a disulfide bond between Cys105 and Cys110 and therefore we termed R-loop (standing for redox-regulated loop), is highlighted in yellow.

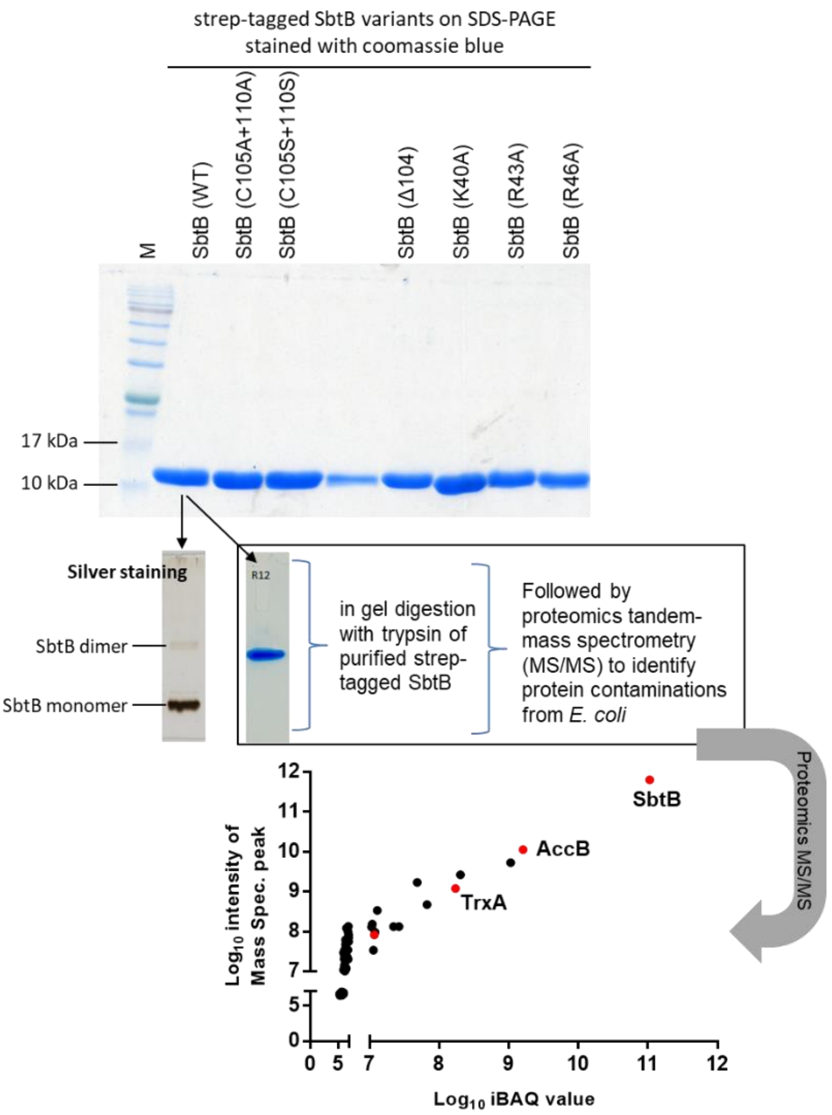


**Fig. S2. SDS-PAGE stained by Coomassie blue for strep-tagged SbtB variants used in this study after purification from *E. coli* expressing the respective protein.** SDS-PAGE showed high degree of purity for all purified SbtB variants. To check for residual protein contaminations from *E. coli*, the wildtype SbtB (WT) was further checked by silver staining, which is more sensitive than Coomassie blue stain, and moreover it was subjected to tandem-mass spectrometry (MS/MS) to identify the *E. coli* proteins, which coeluted with wildtype SbtB (check proteomic dataset associated with this manuscript). The identified proteins were sorted based on iBAQ values of significantly enriched proteins and plotted against the intensity of MS peaks of the identified/defined peptides. The red dots refer to SbtB, thioredoxin-1 (TrxA), glutaredoxin-4 (GrxD), and biotin carboxyl carrier protein of acetyl-CoA carboxylase (AccB). Biotinylated proteins (AccB) are common contaminant of strep-tag purifications. TrxA and GrxD are of special interest as potential targets of SbtB to break the R-loop (check the main text).

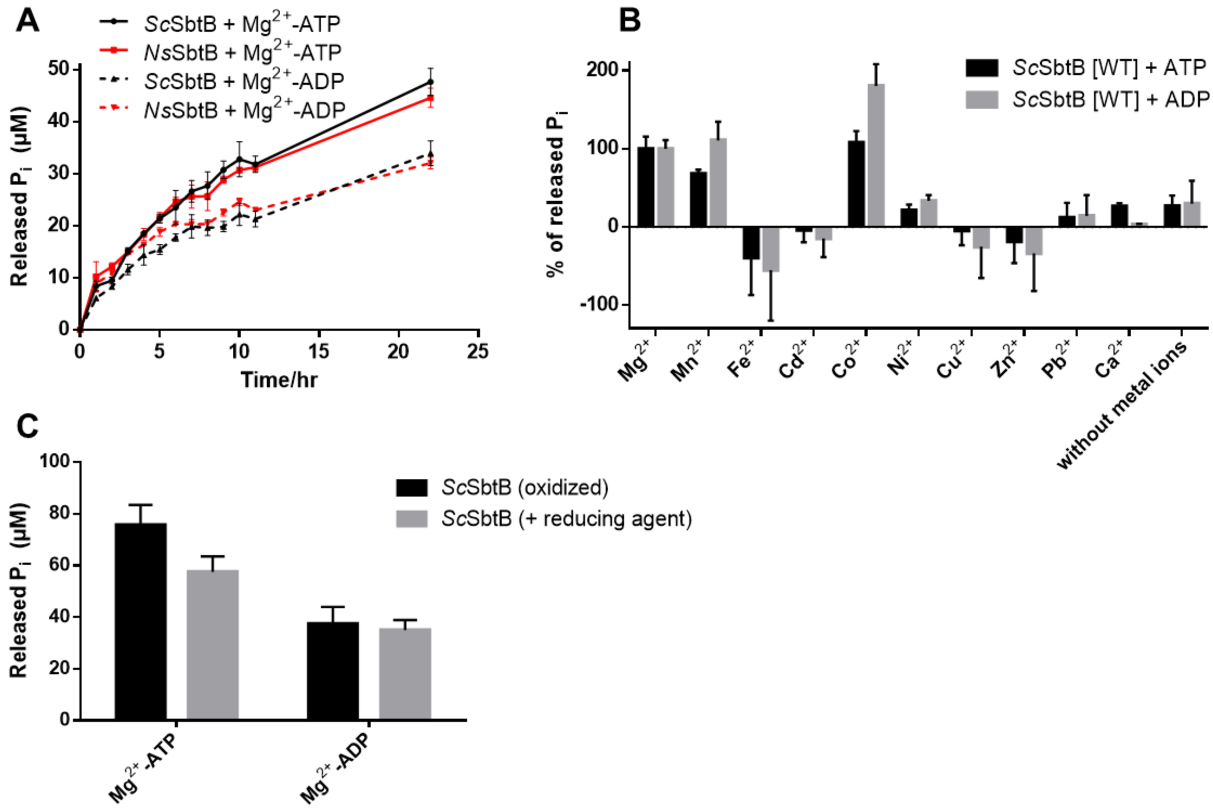


**Fig. S3. Apyrase activity of SbtB proteins via phosphate release assay.** (A) Time course for slow ATP and ADP hydrolysis via *Sc*SbtB and *Ns*SbtB, revealing that ATP and ADP hydrolysis are a common trait among SbtB proteins. The released inorganic phosphate ($P_i$) is shown in µM. (B) Metal influence on *Sc*SbtB apyrase activity, relative to wildtype *Sc*SbtB-activity in presence of $Mg^{2+}$ (100%). The assay was performed in presence of 5 mM of the respective metals. Negative values are indicative of heavily protein precipitation. The assay indicated that $Mn^{2+}$, $Mg^{2+}$ and $Co^{2+}$ could be used as metal ions by *Sc*SbtB. The only metal which can be found in excess inside cells is $Mg^{2+}$, and since high $Co^{2+}$ concentrations is not of physiological relevance, therefore we concluded that $Mg^{2+}$ is most likely the metal used by *Sc*SbtB. (C) Influence of reducing agent on *Sc*SbtB apyrase activity compared to *Sc*SbtB under oxidizing conditions, showing that addition of 1 mM TECP does not influence on *Sc*SbtB apyrase activity.

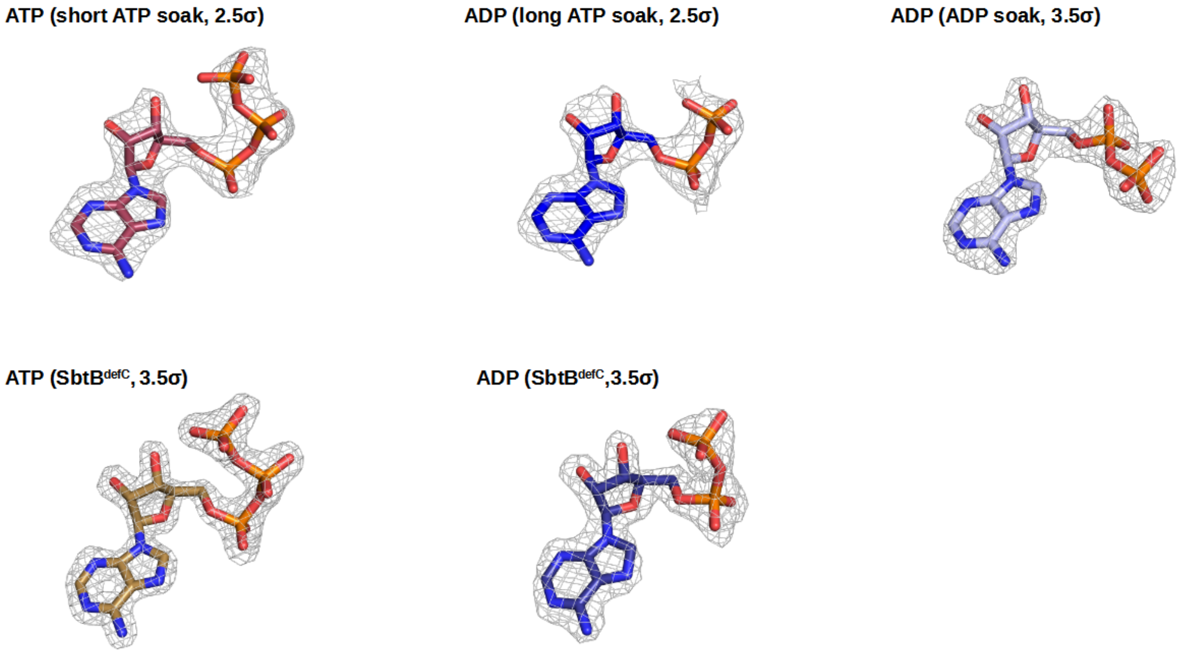


**Fig. S4. Electron densities of nucleotides.**

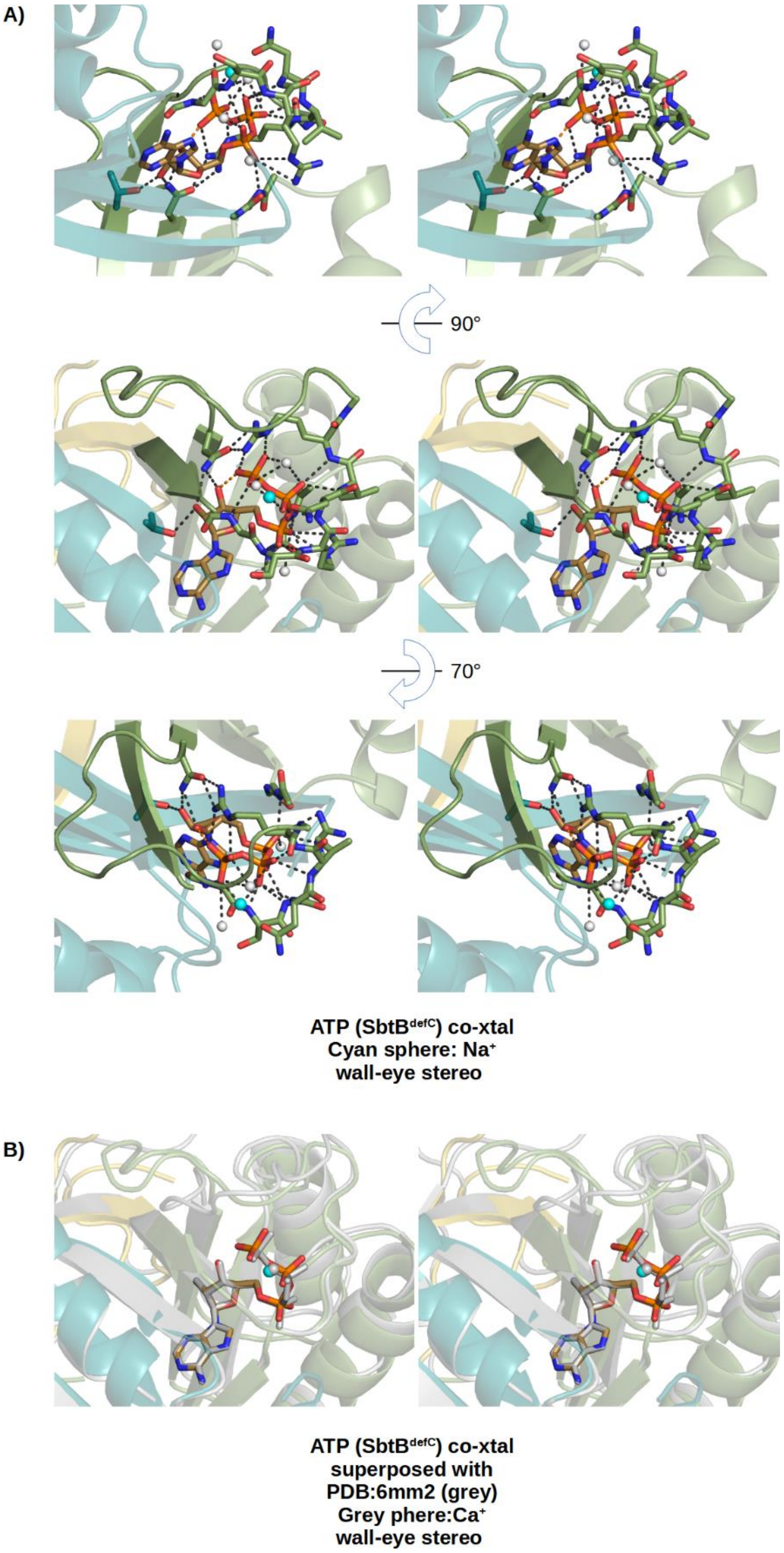


**Fig. S5. Stereo views of the SbtB$^{defR}$:ATP complex.** A) The ATP binding mode is shown in the same orientation as in (Fig. 3), plus two additional orientations, in stereo. B) Stereo superposition of the SbtB$^{defR}$:ATP complex to the SbtB:ATP complex from *Cyanobium* sp. PCC7001 (PDB: 6MM2).

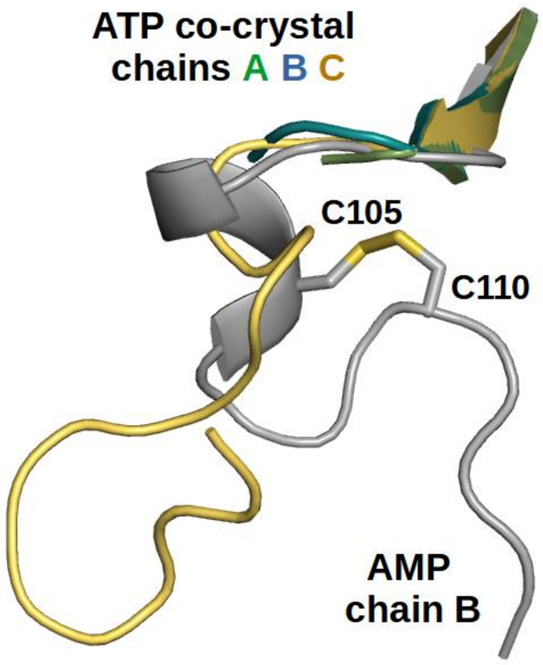


**Fig. S6. Comparison of folded and unfolded R-loop.** The R-loops of the three chains of the SbtB$^{defR}$:ATP co-crystal structure, in which the two R-loop cysteines were substituted by alanine to mimick the reduced state, are superimposed to the oxidized R-loop in the SbtB:AMP co-crystal structure. Obviously, the fold of the oxidized state is not assumed without the disulfide bond and the R-loop completely disordered in two of the three chains.

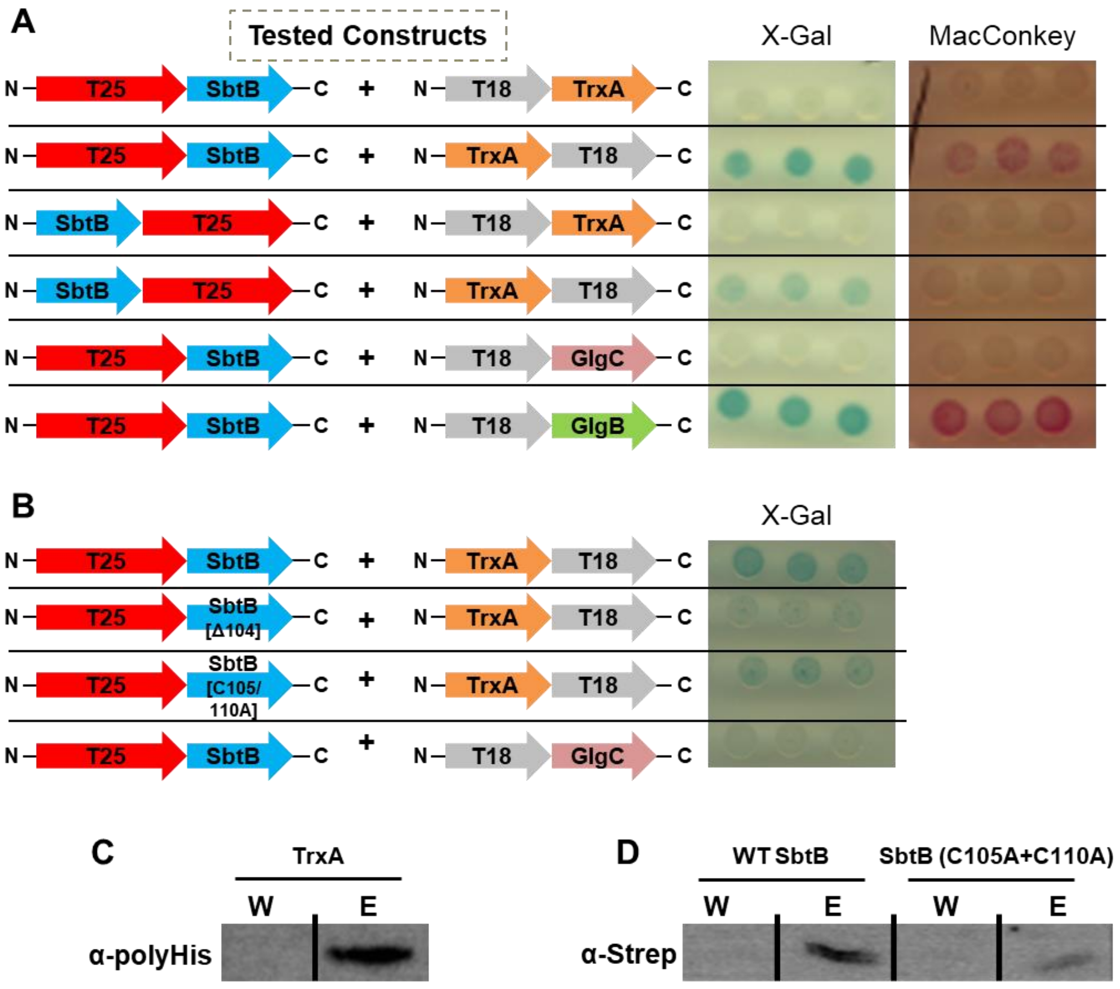


**Fig. S7. Analysis of the interaction between SbtB and TrxA via bacterial two hybrid assay (BACTH) and pulldown assays.** (A) The BACTH assay was performed using *E. coli* cells expressing either N- or C-terminal fusion of Cya-T25 subunit to SbtB together with either N- or C-terminal fusion of Cya-T18 subunit to TrxA, as indicated, on X-Gal or MacConkey reporter plats. N-terminal fusion of Cya-T25 subunit to SbtB together with N-terminal fusion of Cya-T18 subunit with either GlgB or GlgC, was used as positive and negative control, respectively (13). (B) Influence of mutating SbtB R-loop residues on TrxA interaction. The BACTH assay was performed using *E. coli* cells expressing N-terminal fusion of Cya-T18 subunit with TrxA together with N-terminal fusion of Cya-T28 subunit of either wildtype SbtB, or SbtB-(Δ104), or SbtB-(C105A+C110A) as indicated, on X-Gal reporter plat. Positive interaction is evidenced by appearance of a blue or red color on X-Gal or MacConkey reporter plates, respectively. The assay was done using 3-independent/freshly transformed *E. coli* cells, for at least three times to ensure reproducibility. (C and D) Immunoblot blot analysis of SbtB and TrxA interaction in last wash (W) and elution (M) fractions. (C) Strep-tagged SbtB was immobilized and the coelution of TrxA was checked using α-polyHis antibody. (D) His-tagged TrxA was immobilized on $Ni^{2+}$-NTA and the coelution of wildtype SbtB or its variant (C105A+C110A) was checked using α-strep antibody.

### Table S1. Primers and Plasmids

| Primers/ amplification | Sequence (5'→3') | Note/ Ref. |
|---|---|---|
| **Recombinant proteins** | | |
| C-terminal StrepII-tagged *Sc*SbtB (*slr1513*); (pASK-IBA3_*Sc*SbtB-strep plasmid) | 1256_Fw:<br>GTGAAATGAATAGTTCGACAAAAATCTAGATAACGAGGGCAAAAAATG<br>GCTAAACCAGCGAACAAGCTCG | (Selim et al. 2018) |
| | 1257_Rv:<br>AAGCTTATTATTTTTCGAACTGCGGGTGGCTCCAAGCGCTACAGCCCT<br>CAGGGCCACAGAAAG | (Selim et al. 2018) |
| C-terminal StrepII-tagged *Ns*SbtB (*all2133*); (pASK-IBA3_*Ns*SbtB-strep plasmid) | 1664_Fw_all2133_CT strep:<br>GTGAAATGAATAGTTCGACAAAAATCTAGATAACGAGGGCAAAAAATGGCCAAGCCAGCCAAAAAG | (Selim et al. 2021a) |
| | 1665_Rv_all2133_CT strep:<br>CAAGCTTATTATTTTTCGAACTGCGGGTGGCTCCAAGCGCTACAGCCGTCTGGTCCGC | (Selim et al. 2021a) |
| C-terminal StrepII-tagged *Sc*SbtB-Δ104 (pASK-IBA3_*Sc*SbtB-Δ104) | 1256_Fw:<br>GTGAAATGAATAGTTCGACAAAAATCTAGATAACGAGGGCAAAAAATG<br>GCTAAACCAGCGAACAAGCTCG | This study |
| | 1663_Rv_SbtB delta 104:<br>CAAGCTTATTATTTTTCGAACTGCGGGTGGCTCCAAGCGCTGAAAGTATGCCCATAAAGTACTTCTGC | This study |
| C-terminal StrepII-tagged *Sc*SbtB-C105S+C110S (pASK-IBA3_*Sc*SbtB-C105S+C110S) | 1256_Fw:<br>GTGAAATGAATAGTTCGACAAAAATCTAGATAACGAGGGCAAAAAATG<br>GCTAAACCAGCGAACAAGCTCG | This study |
| | 1761_Rv_SbtB-C105+110S<br>CAAGCTTATTATTTTTCGAACTGCGGGTGGCTCCAA-<br>GCGCTGCTGCCCTCAGGGCCGCTGAAAGTATGCCCATAAAGTACTTCTGC | This study |
| C-terminal StrepII-tagged *Sc*SbtB-C105A+C110A (pASK-IBA3_*Sc*SbtB-C105A+C110A) | 1256_Fw:<br>GTGAAATGAATAGTTCGACAAAAATCTAGATAACGAGGGCAAAAAATG<br>GCTAAACCAGCGAACAAGCTCG | This study |
| | 1759_Rv_SbtB-C105+110A<br>CAAGCTTATTATTTTTCGAACTGCGGGTGGCTCCAAGCGCTT-<br>GCGCCCTCAGGGCCTGCGAAAGTATGCCCATAAAGTACTTCTGC | This study |
| C-terminal StrepII-tagged *Sc*SbtB-R46A (pASK-IBA3_*Sc*SbtB-R46A) | 1855_Fw_ R46A_SbtB:<br>AATACCGGTGGCAAGGGTAGCCGTAACGTGGCCTCGTCGGGTCAAC | This study |
| | 1856_Rv_K40,R43,R46:A_SbtB:<br>CATTACCGTGTATCCTTTGGCACCGGATTCG | This study |
| C-terminal StrepII-tagged *Sc*SbtB-R43A (pASK-IBA3_*Sc*SbtB-R43A) | 1854_Fw_R43A_SbtB:<br>AATACCGGTGGCAAGGGTAGCGCCAACGTGCGCTCG | This study |
| | 1856_Rv_K40,R43,R46:A_SbtB:<br>CATTACCGTGTATCCTTTGGCACCGGATTCG | This study |
| C-terminal StrepII-tagged *Sc*SbtB-K40A (pASK-IBA3_*Sc*SbtB-K40A) | 1853_Fw_K40A_SbtB:<br>AATACCGGTGGCGCTGGTAGCCGTAAC | This study |
| | 1856_Rv_K40,R43,R46:A_SbtB:<br>CATTACCGTGTATCCTTTGGCACCGGATTCG | This study |
| N-terminal $His_6$-tagged TrxA (*slr0623*); (pET15b_TrxA-$His_6$ plasmid) | pET15b_slr0623_fw:<br>CAGCAGCGGCCTGGTGCCGCGCGGCAGCCATATGCTCGAGATGAGTGCTACCCCTCAAGTTTC | This study |
| | pET15b_slr0623_rev:<br>CCCTCAAGACCCGTTTAGAGGCCCCAAGGGGTTATGCTAGTTATTGCTCAGCGGTGGCAG-<br>CAGCCAAC | This study |

| BACTH constructs | | |
|---|---|---|
| SbtB-N-terminally tagged with T25 subunit of Cya (pKT25_SbtB_N plasmid) | pKT25_sbtb_fw: CGATTACCTGGCGCGCACGCGGCGGGCTGCAGGGTCGACTATGGCTAAACCAGCGAACAAG | (Selim et al. 2021a) |
| | pKT25_sbtb_rev: GGCCGAATTCTTAGTTACTTAGGTACCCGGGGATCCTCTAGTTAACAGCCCTCAGGGCCAC | (Selim et al. 2021a) |
| SbtB-C-terminally tagged with T25 subunit of Cya (pKT25_SbtB_C plasmid) | Fw: GAATTGTGAGCGGATAACAATTTCACACAGGAAACAGCTATGATGGCTAAACCAGCGAAC | This study |
| | Rv: CGGCGTTTGCGTAACCAGCCTGATGCGATTGCTGCATGGTACAGCCCTCAGGGCCACAGAAAG | This study |
| SbtB-N-terminally tagged with T25 subunit of Cya (pKT25_SbtB_[C105A+C110A] plasmid) | Fw: TTCGGTGACCGATTACCTGGCGCGCACGCGGCGGGCTGCAGCTAAACCAGCGAACAAGCTC | This study |
| | Rv: ACGACGGCCGAATTCTTAGTTACTTAGGTACCCGGGGATCTTATGCGCCCTCAGGGCCTGC | This study |
| SbtB-N-terminally tagged with T25 subunit of Cya (pKT25_SbtB_[Δ104] plasmid) | Fw: TTCGGTGACCGATTACCTGGCGCGCACGCGGCGGGCTGCAGCTAAACCAGCGAACAAGCTC | This study |
| | Rv: ACGACGGCCGAATTCTTAGTTACTTAGGTACCCGGGGATCTTAGAAAGTATGCCCATAAAGT | This study |
| TrxA-N-terminally tagged with T18 subunit of Cya (*slr0623*_N); (pUT18_TrxA_N plasmid) | Fw: GCGGCGGCCGTCGCTGGGCGCAGTGGAACGCCACTGCAGGAGTGCTACCCCTCAAGTTTC | This study |
| | Rv: TTAGTTATATCGATGAATTCGAGCTCGGTACCCGGGGATCTTAAAGA-TATTTTTCTAGGGTGCTGG | This study |
| TrxA-C-terminally tagged with T18 subunit of Cya (*slr0623*_C); (pUT18_TrxA_C plasmid) | Fw: CAATTTCACACAGGAAACAGCTATGACCATGATTACGCCAATGAGTGCTACCCCTCAAGT | This study |
| | Rv: CTGAATTCGAGCTCGGTACCCGGGGATCCTCTAGAGTCGAAAGATATTTTTCTAGGGTGCTGGC | This study |
| GlgB-N-terminally tagged with T18 subunit of Cya (*sll0158*_N); (pUT18_GlgB_N plasmid) | pUT18 glgB sll0158 fw GACCATGATTACGCCAAGCTTGCATGCCTGCAGGTCGACTATGACCTACACCATCAACG | (Selim et al. 2021a) |
| | pUT18 glgB sll0158 rev CCTCGCTGGCGGCTGAATTCGAGCTCGGTACCCGGGGATCAGCTATGTTGCTAGCCTCTTC | (Selim et al. 2021a) |
| GlgB-C-terminally tagged with T18 subunit of Cya (*sll0158*_C); (pUT18_GlgB_C plasmid) | pUT18c glgB sll0158 fw GCCGTCGCTGGGCGCAGTGGAACGCCACTGCAGGTCGACTATGACCTACACCATCAACG | (Selim et al. 2021a) |
| | pUT18c glgB sll0158 rev TTAGTTATATCGATGAATTCGAGCTCGGTACCCGGGGATCAGCTATGTTGCTAGCCTCTTC | (Selim et al. 2021a) |
| GlgC-C-terminally tagged with T18 subunit of Cya (*slr1176*_C); (pUT18_GlgC_C plasmid) | pUT18 glgC AGP slr1176 fw: GACCATGATTACGCCAAGCTTGCATGCCTGCAGGTCGACTGTGTGTTGTTGGCAATCGAG | This study |
| | pUT18 glgC AGP slr1176 rev: CCTCGCTGGCGGCTGAATTCGAGCTCGGTACCCGGGGATCGATTACCGTGCCGTCGGCGATC | This study |
| GlgC-N-terminally tagged with T18 subunit of Cya (*slr1176*_N); (pUT18_GlgC_N plasmid) | pUT18c glgC AGP slr1176 fw: GCCGTCGCTGGGCGCAGTGGAACGCCACTGCAGGTCGACTGTGTGTTGTTGGCAATCGAG | This study |
| | pUT18c glgC AGP slr1176 rev: TTAGTTATATCGATGAATTCGAGCTCGGTACCCGGGGATCGATTACCGTGCCGTCGGCGATC | This study |

### Table S2. Data Collection and Refinement Statistics

| Structure | SbtB, short ATP soak | SbtB, long ATP soak | SbtB, ADP soak | SbtB C105A+C110A variant, ATP co-crystal | SbtB (Δ104) variant, ADP co-crystal |
|---|---|---|---|---|---|
| **PDB code** | **7R2Y** | **7R2Z** | **7R30** | **7R31** | **7R32** |
| **Data collection** | | | | | |
| Space group | P3$_2$ | P3$_2$ | P3$_2$ | P4$_1$ | P4$_1$ |
| Cell parameters | a = b = 63.83 *Å*, c = 81.23 *Å* | a = b = 60.80 *Å*, c = 78.44 *Å* | a = b = 63.64 *Å*, c = 82.21 *Å* | a = b = 73.19 *Å*, c = 89.01 *Å* | a = b = 73.98 *Å*, c = 88.42 *Å* |
| Wavelength (*Å*) | 1.000 | 1.000 | 1.000 | 1.000 | 1.000 |
| Resolution limits (*Å*)[a] | 32.73-2.15 (2.25-2.15) | 31.45-2.40 (2.54-2.40) | 32.95-1.90 (2.01-1.90) | 33.85-1.52 (1.61-1.52) | 56.74-1.75 (1.85-1.75) |
| Unique reflections | 20143 (3177) | 12684 (2032) | 29447 (4769) | 71568 (11257) | 47404 (7186) |
| Completeness (%) | 99.6 (97.6) | 99.8 (98.8) | 100 (100) | 98.8 (96.8) | 98.8 (97.7) |
| Redundancy | 10.2 (9.51) | 7.23 (6.79) | 10.3 (10.3) | 11.9 (8.11) | 9.89 (10.1) |
| I/σI | 27.0 (1.70) | 20.8 (2.28) | 20.5 (1.58) | 24.4 (1.73) | 17.0 (2.03) |
| $R_{merge}$ (%) | 4.4 (139.5) | 5.8 (78.4) | 5.8 (153.7) | 5.4 (109.4) | 8.5 (142.2) |
| CC(1/2) | 100 (75.1) | 99.9 (86.3) | 99.9 (71.3) | 99.9 (93.2) | 99.9 (74.2) |
| **Refinement** | | | | | |
| Resolution limits (*Å*) | 32.73-2.15 (2.20-2.15) | 31.45-2.40 (2.46-2.40) | 32.95-1.90 (1.95-1.90) | 33.85-1.52 (1.56-1.52) | 56.74-1.75 (1.80-1.75) |
| $R_{cryst}$ (%) | 18.0 (36.6) | 18.6 (45.4) | 17.1 (36.1) | 18.8 (39.1) | 18.6 (35.8) |
| $R_{free}$ (%) | 21.2 (43.9) | 21.6 (47.7) | 19.5 (34.2) | 20.6 (36.9) | 20.9 (37.8) |
| Protein molecules / asymmetric unit | 3 | 3 | 3 | 3 | 3 |
| Mean B value ($Å^2$) | 70.2 | 66.2 | 51.0 | 28.1 | 33.2 |
| **Ramachandran Statistics**[b] | | | | | |
| Core regions (%) | 93.5 | 93.3 | 94.9 | 96.2 | 95.8 |
| Allowed regions (%) | 99.2 | 99.6 | 100 | 100 | 99.6 |

[a] Values in parenthesis refer to the highest-resolution shell.

[b] Ramachandran statistics were determined with PROCHECK.
